# C5aR1 antagonism suppresses inflammatory glial gene expression and alters cellular signaling in an aggressive Alzheimer’s model

**DOI:** 10.1101/2023.08.22.554306

**Authors:** Nicole D. Schartz, Heidi Y. Liang, Klebea Carvalho, Shu-Hui Chu, Adrian Mendoza-Arvilla, Tiffany J. Petrisko, Angela Gomez-Arboledas, Ali Mortazavi, Andrea J. Tenner

## Abstract

Alzheimer’s disease (AD) is the leading cause of dementia in older adults, and the need for effective, sustainable therapeutic targets is imperative. Pharmacologic inhibition of C5aR1 reduces plaque load, gliosis and memory deficits in animal models. However, the cellular basis underlying this neuroprotection and which processes were the consequence of amyloid reduction vs alteration of the response to amyloid were unclear. In the Arctic model, the C5aR1 antagonist PMX205 did not reduce plaque load, but deficits in short-term memory in female mice were prevented. Hippocampal single cell and single nucleus RNA-seq clusters revealed C5aR1 dependent and independent gene expression and cell-cell communication. Microglial clusters containing neurotoxic disease-associated microglial genes were robustly upregulated in Arctic mice and drastically reduced with PMX205 treatment, while genes in microglia clusters that were overrepresented in the Arctic-PMX205 vs Arctic group were associated with synapse organization and transmission and learning. PMX205 treatment also reduced some A-1 astrocyte genes. In spite of changes in transcript levels, overall protein levels of some reactive glial markers were relatively unchanged by C5aR1 antagonism, as were clusters associated with protective responses to injury. C5aR1 inhibition promoted signaling pathways associated with cell growth and repair, such as TGFβ and FGF, in Arctic mice, while suppressing inflammatory pathways including PROS, Pecam1, and EPHA. In conclusion, pharmacologic C5aR1 inhibition prevents cognitive loss, limits microglial polarization to a detrimental inflammatory state and permits neuroprotective responses, as well as leaving protective functions of complement intact, making C5aR1 antagonism an attractive therapeutic strategy for individuals with AD.

**One Sentence Summary:** Pharmacologic inhibition of C5aR1 suppresses disease-enhancing processes and promotes disease mitigating pathways in an aggressive model of Alzheimer’s disease.

## INTRODUCTION

Alzheimer’s disease (AD) is the most common form of dementia in the elderly, with an estimated 6.5 million people currently diagnosed in the US, with an expected doubling by 2060 (*1*). AD pathology is characterized by the accumulation of extracellular amyloid beta (Aβ) plaques, hyperphosphorylated tau, and neuronal loss ultimately resulting in cognitive decline (*2*). Although Aβ deposition and tau phosphorylation are the hallmarks of AD, evidence suggests that inflammation triggered by these events may contribute to onset and acceleration of functional loss and may be a more appropriate therapeutic target to prevent disease progression (*3*).

Complement activation is critical for the rapid recognition and clearance of pathogens, apoptotic cells, and cellular debris. Activation of the complement system in the CNS promotes opsonization via C3b/iC3b, myeloid cell recruitment via the production of C3a and C5a, and cell lysis via the membrane attack complex (MAC) (*4*). Classical complement pathway component C1q has been largely shown to be neuroprotective in *in vitro* systems by clearing apoptotic cells, modulating subsequent inflammatory cytokine production and directly enhancing survival of neurons (*5–8*). However, under pathological conditions such as AD, additional complement components that enable the activation of the entire complement cascade may also be expressed (*9*), leading to cleavage of C5 into C5a and C5b fragments resulting in induction of inflammation as well as neuronal damage. In human AD and mouse models of AD, complement components C1q, C3b, and C4b co-localize with fibrillar Aβ plaques ((*10–13*) and reviewed in (*14*)) providing evidence of pathway activation.

The primary C5a receptor (C5aR1, aka CD88) is expressed at low levels in microglia in the CNS under physiological conditions, but under pathological conditions it can be highly upregulated and may be expressed by other cell types such as neurons and endothelial cells (*15*). C5aR1 is most highly expressed in plaque-associated microglia (*16*), and genetic ablation or inhibition of C5aR1 or C5a in AD mouse models restores cognitive performance and reduces neuroinflammation (*17–21*). C5a-C5aR1 engagement results in potent pro-inflammatory responses via MAPK signaling, resulting in generation of pro-inflammatory cytokines, and is known to synergize with toll-like receptors to enhance pro-inflammatory cytokine responses (*22*). C5aR1 exerts its effects via cAMP and ERK1/2 signaling as well as recruitment of β-arrestin 2 (*23*). Thus, the neuroprotective effect resulting from C5aR1 inhibition may be due to intercellular signaling pathways involving microglia and/or other cell types including neurons, astrocytes, oligodendrocytes, or endothelial cells.

PMX205 is a cyclic hexapeptide noncompetitive inhibitor of C5aR1 (*24*). With high oral bioavailability, low accumulation in the blood, brain, or spinal cord, and efficacy at crossing the blood brain barrier, it is a viable candidate for chronic treatment to block C5a-C5aR1 signaling in the brain (*24, 25*). PMX205 treatment has been reported to exert neuroprotective effects in models of neurodegeneration, including experimental autoimmune encephalomyelitis (*26*), amyotrophic lateral sclerosis (*27*), and spinal cord injury (*28*). We have previously reported that PMX205 treatment initiated at the onset of amyloid deposition in the Tg2576 mouse model reduces amyloid plaque accumulation and dystrophic neurites, alters gene expression in microglia, and improves cognitive performance (*20, 21*). In the present study, we treated the aggressive Arctic48 mouse model of AD with PMX205 after substantial amyloid plaques were already present to determine the protective effects of C5aR1 inhibition in advanced amyloid pathology. To this end, we tested spatial memory performance with the Y maze and assessed microglial or nucleus preparations by RNA-seq. While Arctic mice had a robust upregulation of inflammatory microglial gene expression, PMX205 treatment suppressed intercellular signaling in the Arctic brain, reduced inflammatory microglial gene expression, and promoted protective signaling pathways from astrocytes and microglia including transforming growth factor-β (TGFb), bone morphogenetic protein (BMP), and fibroblast growth factor (FGF). Spatial memory deficits were observed in Arctic females, not males at 10 mo of age, suggesting sex-specific progression of disease pathology. However, treatment with PMX205 was sufficient to prevent short-term spatial memory loss in Arctic female mice.

## RESULTS

### Treatment with PMX205 suppresses complement gene transcription in Arctic mice

Mice were treated with PMX205 or vehicle from ages 7.5 to 10 months, when plaque accumulation in the hippocampus rises and plateaus (*19*) (**Figure 1A**). Mice did not lose weight during the course of the treatment (**Figure S1A-B**) and consumed an average of 6-10 mL of water or PMX205 in water per day (**Figure S1C-D**), resulting in an average dose of 4.76 mg/kg and 5.36 mg/kg of PMX205 per day for WT and Arctic (Arc) mice, respectively (**Figure S1E**), consistent with doses in previous studies (*21*). WT females had a higher average daily dose of PMX205 than WT males due to the higher body weight of WT males (**Figure S1F**).

**Fig.1:**
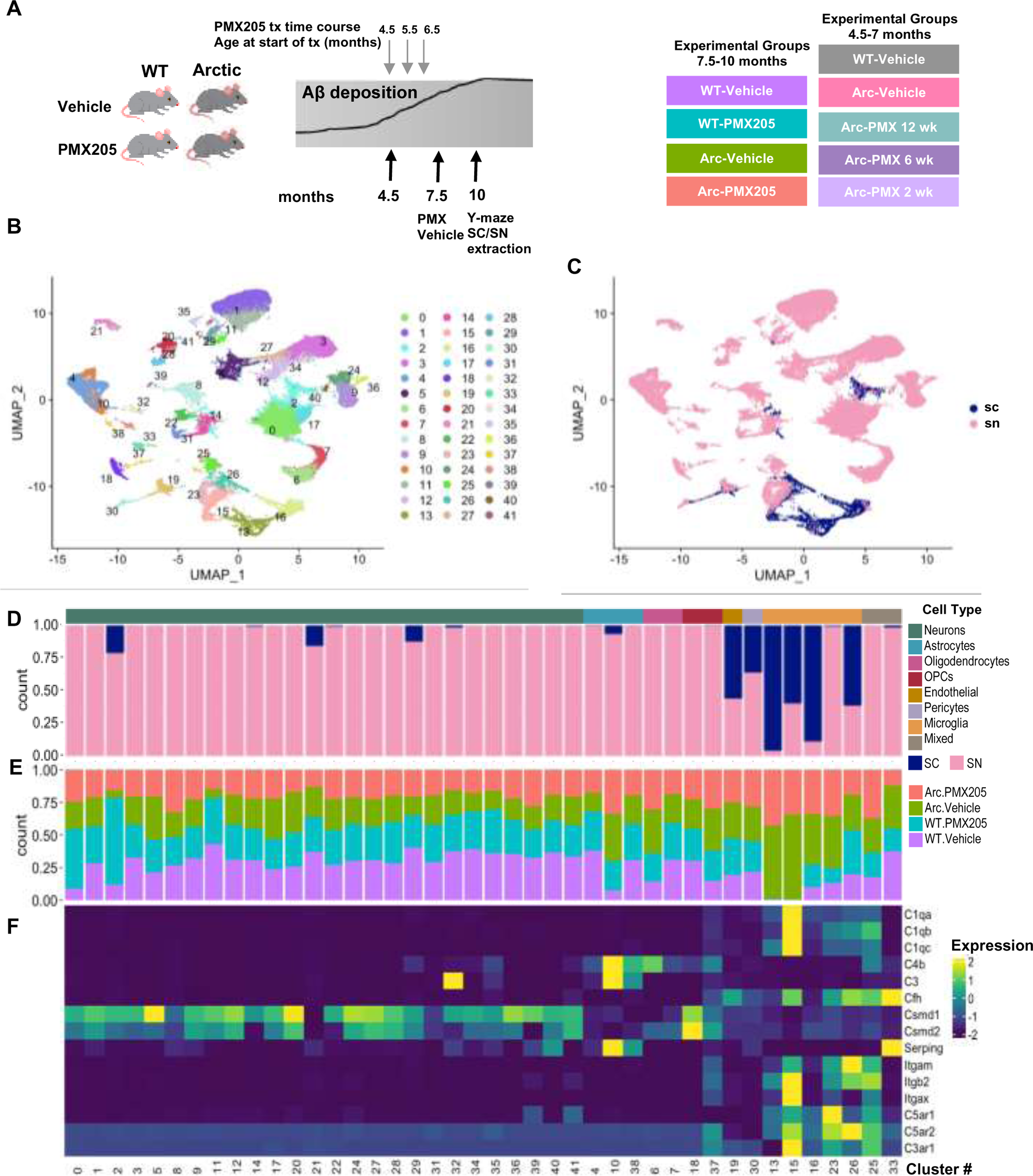
Clustering of transcriptome from single nucleus and single cells in wild type and Arctic mice. (**A**) Experimental timeline for PMX205 treatments (**B**) Seurat cluster identification of combined cells/nuclei transcriptome. (**C**) Seurat cluster identification of cells derived from single cells (SC) or single nucleus (SN) sequencing. (**D**) Proportion of cells in each cluster originating from SC or SN transcriptome. (**E**) Proportion of cells in each cluster by treatment/genotype. (**F**) Expression of complement pathway components, receptors or regulators by cell type.

Transcripts from single cell microglia and single nucleus libraries derived from hippocampi were combined and clusters were identified with Seurat. 42 clusters containing 54,157 cells/nuclei were identified (**Figures 1B-C, 2A**). The majority of clusters were derived from single nucleus populations, while microglial clusters were comprised of mostly single cells although one microglial cluster was exclusively derived from single nucleus (**Figure 1D**). Interestingly, some microglial clusters (13, 16 and 23) from SN- and SC-RNAseq were segregated between nuclear and whole cell locations, which is likely the result of compartment-dependent differences in mRNA half-life and/or populations (**Figure 1C-D**). In addition, we identified specific clusters that were over-represented in the Arc samples compared to WT, including microglial clusters 13 and 15, and astrocyte cluster 10 (**Figure 1E**). Within these clusters, microglia clusters, particularly those unique to Arctic mice, expressed the highest levels of complement genes, while neuronal populations expressed relatively high levels of regulatory complement genes Csmd1 and Csmd2 (**Figure 1F**). The astrocyte cluster that was enriched in Arc samples expressed high levels of C3, C4b, and Serping1.

Of the 42 Seurat clusters, 26 were identified as neurons, 5 as microglia, and 3 as astrocytes (**Figure 1B, 1D**). Among the single nucleus samples, neurons made up the majority of cell types sequenced from each of the treatment groups. We also identified 2 clusters of oligodendrocytes and 2 of oligodendrocyte progenitor cells (OPCs), as well as single clusters of endothelial cells, pericytes, and two “mixed” clusters that are likely a mixture of astrocytes and microglia (**Figure 1D and 2A, 2B**).

**Fig.2:**
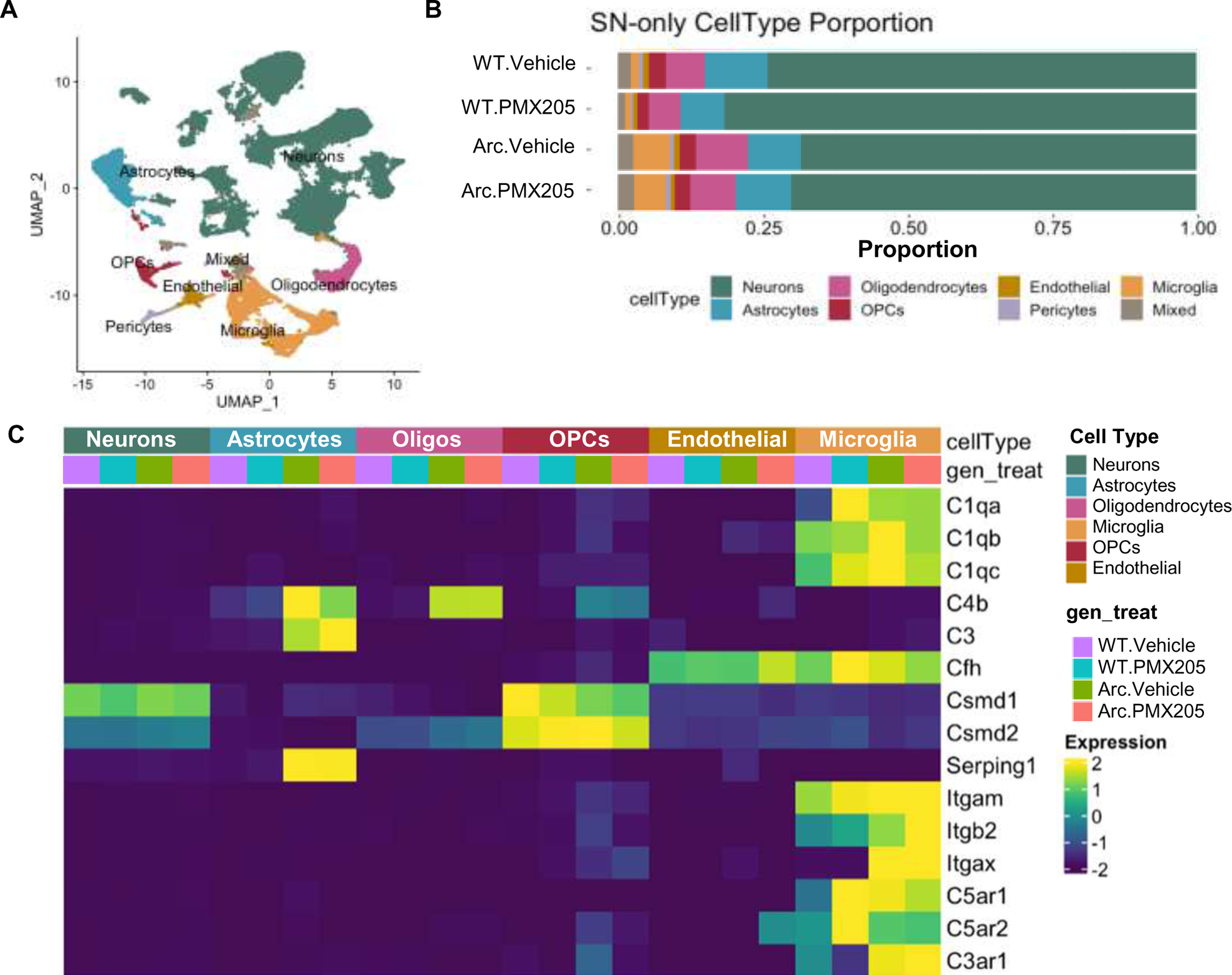
Single cell and single nucleus RNA-Seq clustered by cell type reveals cell-specific complement gene expression. Microglia or nuclei were isolated from hippocampi, fixed, and sequenced. (**A**) U-Map of all cell types with all treatment groups included (counts per cell type were: Neurons 37512, Astrocytes 4612, Oligodendrocytes 3478, Microglia 4891, OPCs 1259, Endothelial 898, Mixed 1017, Pericytes 490). (**B**) Proportion of isolated cell types in the single nucleus RNA-seq by genotype and treatment group. (**C**) Differential cell type-specific expression of complement pathway components and regulators of the complement system derived from single nucleus RNA-seq.

Because we used a treatment to modulate complement activation-mediated signaling, we further explored the relative complement gene expression in each cell cluster and genotype/treatment group (**Figure S2A**). Neuronal clusters expressed relatively high levels of complement regulatory genes Csmd1 and Csmd2 (irrespective of genotype), and astrocytes had high transcript levels of C4b, C3 and Serping1 (C1 Inhibitor) expressed predominantly in Arc mice (**Figure 2C**). Microglial clusters expressed C1qa, C1qb, C1qc, Cfh (Factor H), Itgam (CD11b), Itgb2 (CD18), Itgax (CD11c), C5ar1, C5ar2, and C3ar1 (**Figure 2C and 1F**). When only looking at SN samples, neuronal, endothelial, and OPC expression of complement genes was unchanged, while complement genes in astrocytes and microglia were upregulated in Arc compared to WT (**Figure 2C**). To discriminate in a more focused analysis, microglia and astrocyte data were separately re-clustered.

### PMX205 treatment results in downregulation of inflammatory and neurotoxic microglia genes

We subclustered the SC and SN microglia clusters without the other brain cell types to identify 11 clusters (**Figure 3A**) and calculated the proportional representation of each cluster per treatment group (**Figure 3B**). Based on very low expression levels of known microglial genes such as Csf1r, Hxb, Cx3cr1, we determined that cluster M11 had limited microglia, and was excluded from further analysis (**Figure S2B**). Comparisons of the proportional distributions of microglial clusters within treatment groups revealed clusters that were enriched in WT cells (cluster M5), clusters that were specific to Arctic cells regardless of PMX205 treatment (clusters M3 and M8), clusters that were highly enriched in Arctic cells but ablated with treatment of PMX205 (clusters M4, M6 and M9), and clusters that were enriched in cells upon PMX205 treatment (cluster M1) (**Figure 3B**). Hierarchical clustering was used to segregate microglial clusters based on similarities of marker gene expression and then to assess proportions among SC/SN preps (**Figure 3C**) and treatment groups (**Figure 3D**).

**Fig.3:**
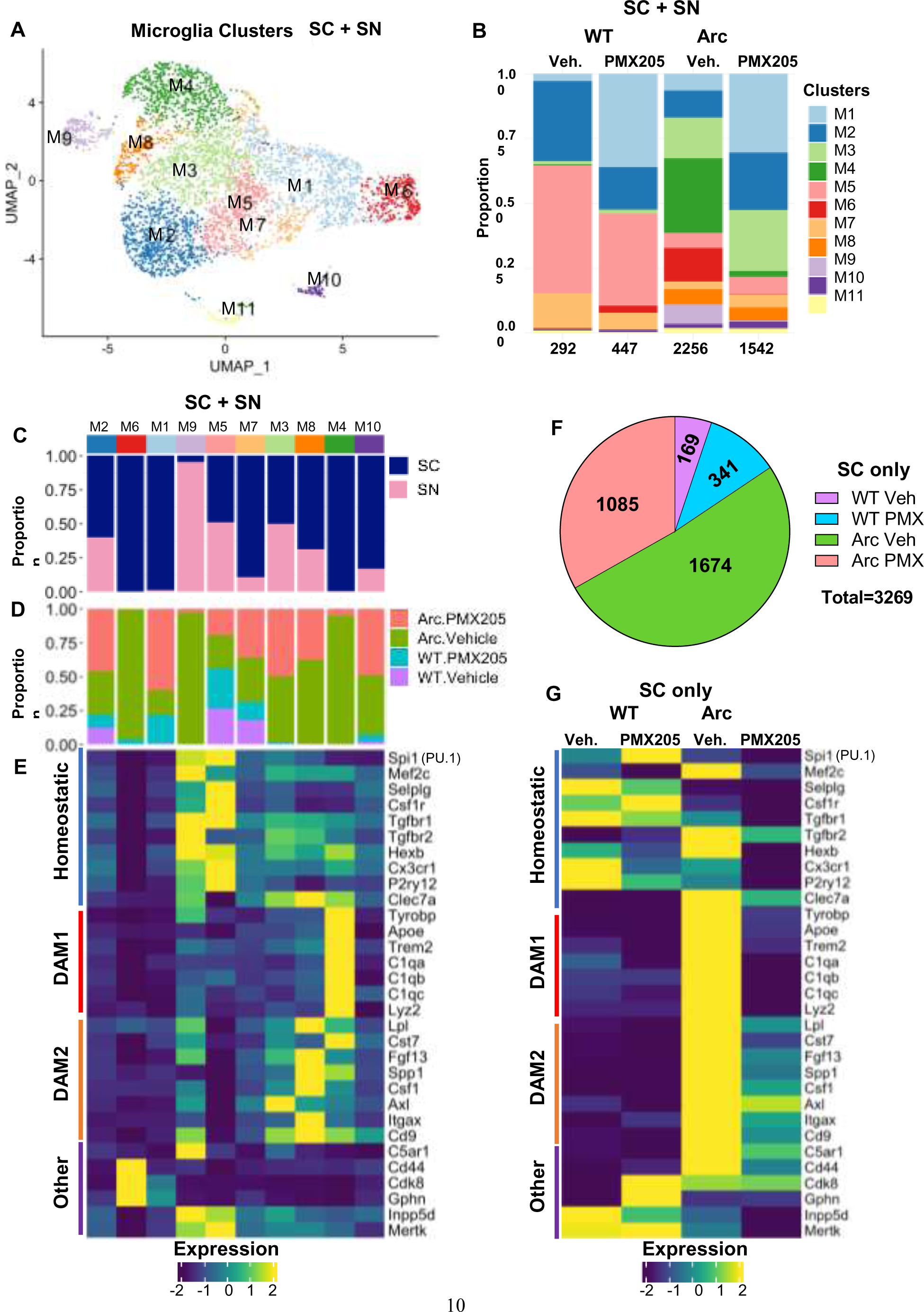
DAM1 gene expression is suppressed in Arctic-PMX205 hippocampal microglia. (**A**) Cells/nuclei identified as microglia were re-clustered separately. (**B**) Proportion of microglia (SC+SN) clusters in WT-Veh, WT-PMX, Arc-Veh, and Arc-PMX samples. (**C**) Proportion of cells in each cluster originating from SC or SN transcriptome. (**D**) Proportion of cells/nuclei in each cluster by treatment/genotype. (**E**) Relative expression of genes representative of homeostatic microglia, DAM1, DAM2, or other genes of interest within the different microglial clusters. (**F**) Pie chart demonstrating proportion of SC microglia samples derived from different treatment groups. (**G**) Relative expression of homeostatic, DAM1, or DAM2 genes in different treatment groups, with SC data alone.

Based on the current categorization that is widely used in the literature (*29*), we assessed relative expression of homeostatic, DAM1, and DAM2 genes by clusters as well as other genes of interest with SC and SN preparations grouped together (**Figure 3C-E**) and with SC microglia analyzed separately (**Figure 3G**). DAM1 gene transcripts (Cluster M4, **Figure 3C-E**) are primarily found in the single cell, not the nuclear preparations, and interestingly, DAM1 genes were overexpressed in Arc females compared to Arc males (**Figure S3**). Notably, cluster M5, with high expression of homeostatic microglial genes, such as Selplg and P2ry12 (**Figure 3E and Supplemental File 1**), was highly represented in WT microglia, regardless of PMX205 treatment (**Figure 3G**). Clusters M3 (Axl) and M8 (Itgax and Spp1) were present only in the Arc mice regardless of PMX205 treatment and had low DAM1 gene expression. M8 had the highest relative expression of DAM2 genes (**Figure 3E**). This suggests that the DAM2-induced responses to injury represented in clusters M3 and M8 were independent of C5aR1 signaling.

In contrast, clusters M4, M6 and M9 were highly induced in the Arc hippocampus, but only minimally detected in Arc-PMX205. Top genes of cluster M4 included Apoe, C1qa, C1qb, C1qc, Tyrobp, Trem2, Lyz2 and Cst7 (**Figure 3E, Supplemental File 1**). These DAM1 genes were highly expressed in Arc microglia compared to WT microglia and were downregulated in Arc-PMX205 microglia (**Figure 3F**). GO analysis for cluster M4 identified pathways associated with synapse pruning, microglial cell activation, positive regulation of cell death, and regulation of myeloid cell differentiation suggesting that this population may be contributing to detrimental effects that are blocked by inhibition of C5aR1 signaling (**Figure S4A**). Similarly, cluster M9 (with high expression of C5ar1) expressed disease associated microglial genes Inpp5d, Mertk, and Tgfbr2 (**Supplemental File 1**). GO analysis of cluster M9 genes revealed pathways associated with vasculogenesis, cell activation, and fatty acid metabolic processes (**Figure S4B**). Cluster M6 was differentiated from other clusters by only 16 genes (**Supplemental File 1**). Among those genes were modulators of the immune system, Lars2, that supports a higher metabolic state, Cmss1 involved in translation, and Cdk8 that promotes inflammation (*30*). Treatment with PMX205 suppresses those injury-induced inflammation enhancing genes, suggesting that C5aR1 mediates inflammatory injury response to amyloid accumulation in the hippocampus.

In addition, microglia cluster M2 made up a smaller proportion in Arc cells relative to WT, but was largely rescued in Arc-PMX205 microglia. Genes found in microglial cluster M2 were associated with trans-synaptic signaling, synapse organization, regulation of glutamatergic transmission, and learning (**Figure S4C**). These findings suggest that C5aR1 inhibition in Arc mice enhances pathways associated with synaptic plasticity and learning while suppressing pathways associated with glial activation, synaptic loss, and cell death.

Finally, Cluster M1, which is dependent on PMX205 treatment regardless of genotype, contained genes associated with mRNA splicing and microtubule cytoskeleton organization and processes. Although this cluster was not defined by many genes, it may indicate that inhibition of C5aR1 may promote glial structural integrity and neuroprotective functions. In summary, when assessed by treatment group/genotype, homeostatic genes were highly represented in WT-Veh and WT-PMX205 microglia, while DAM1 genes were highly upregulated in Arc-Veh cells. Treatment with PMX205 had a strong effect in reducing DAM1 gene expression, but less so on DAM2 expression. Of note, the microglia that had highest DAM1 gene expression were derived from female Arc-Veh mice and may suggest an accelerated timeline to neuroinflammation in females (**Figure S3**).

### PMX205 treatment reduces expression of some neurotoxic astrocyte genes

Astrocyte clusters were also isolated and re-clustered for further exploration of astrocyte subpopulations. Twelve astrocyte (A) clusters were defined (**Figure 4A**) and confirmed to contain astrocytes based on gene expression (**Figure S2C**). Of these, clusters A1 and A2 were dominant in WT cells, while cluster A3 was specific to Arc cells regardless of PMX205 treatment and cluster A7 was high in Arc cells and ablated with PMX205 treatment (**Figure 4B**). In contrast, cluster A6 was depleted in Arc but rescued in ArcPMX205.

**Fig.4:**
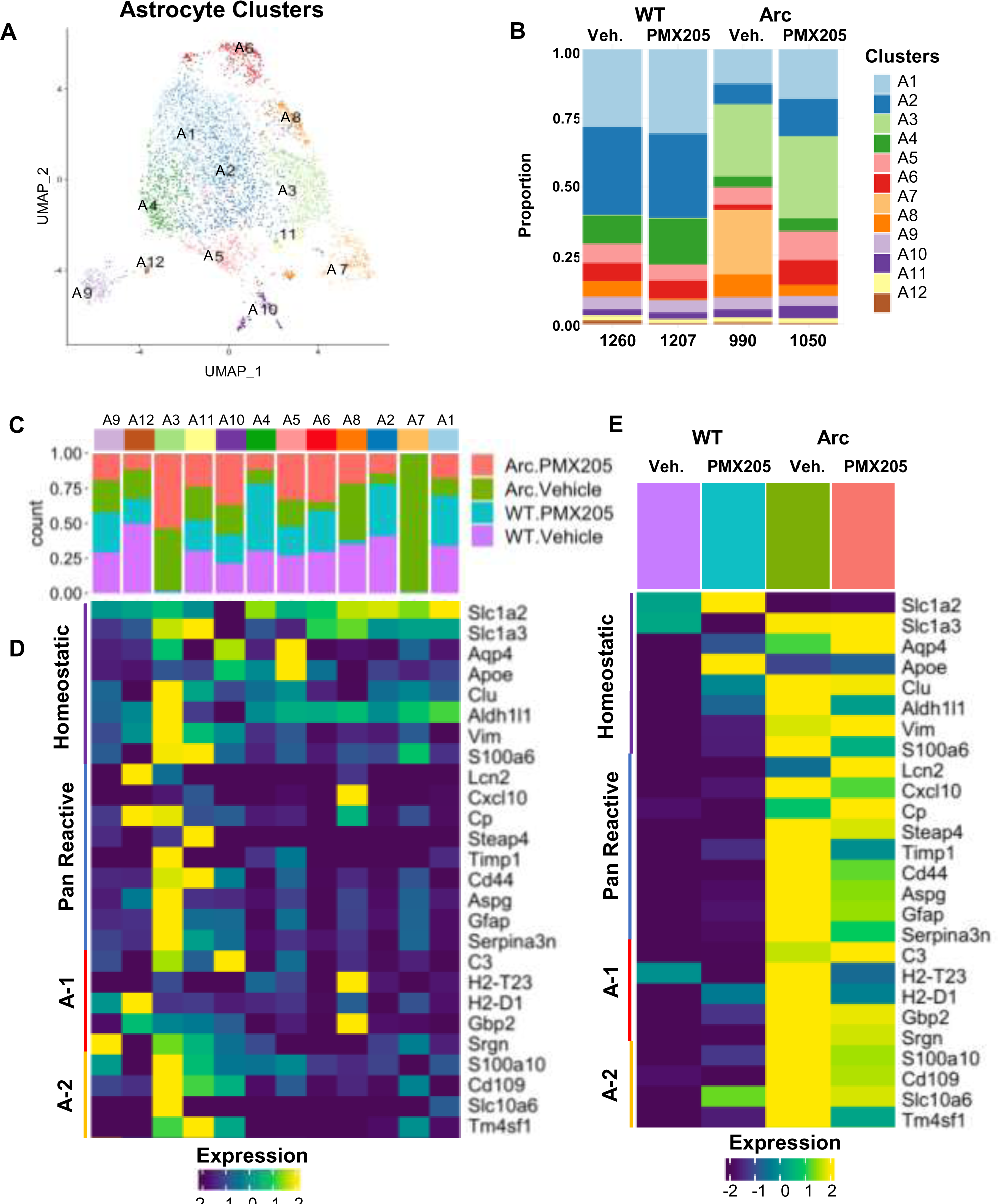
Reactive Astrocyte gene expression is largely suppressed in Arctic-PMX205 hippocampus. (**A**) Cells identified as astrocytes were re-clustered separately. (**B**) Proportion of astrocyte populations in WT-Veh, WT-PMX, Arc-Veh, and Arc-PMX samples. (**C**) Proportion of cells in each cluster by treatment/genotype. (**D**) Relative expression of genes representative of homeostatic astrocytes, pan-reactive, A-1 neurotoxic, or A-2 neuroprotective astrocytes within the different astrocyte clusters. (**E**) Relative expression of homeostatic, pan-reactive, A-1, or A-2 genes in the different treatment groups.

To better understand the role of the Arc-specific astrocyte clusters, we performed GO analysis on the cluster-defining genes. Cluster A3, which included Gfap, C4b and Cd44 amongst its top genes (**Supplemental file 1**) had GO terms associated with tyrosine kinase signaling pathways, nerve development, response to wounding, and JAK-STAT pathway (**Figure S5A**). Interestingly, cluster A3 was highly enriched for pan-reactive and A-2 (*31*) reactive genes (**Figure 4D**), thus supporting that the function of this population is associated with disease-mitigating response to inflammation and injury that is independent of C5aR1 signaling.

Cluster A6 was suppressed in Arc astrocytes and was rescued by PMX205 (**Figure 4B-C**). Cluster A6 GO terms included axon guidance, receptor recycling, protein processing, and neuron differentiation (**Figure S5B**). Interestingly, cluster A10, increased slightly in Arc but more so in PMX205 treated Arc, had GO terms including defense response to virus, cellular response to interferon β, and regulation of response to cytokine stimulus (**Figure S5C**). The enhancement of cluster A10 in Arc-PMX205 suggests that C5aR1 inhibition in Arc mice enables a protective response from astrocytes in response to perceived injury. Thus, enhancement of clusters A6 and A10 by PMX205 is consistent with a heightened phenotype in astrocytes when C5a-C5aR1 signaling is inhibited that results in support of neuronal functions in response to inflammatory stimulus.

Cluster A7, which was unique to Arc astrocytes and not found in Arc-PMX205 cells, was only defined by 10 genes including Arhgef4, Rgs6, and Enox1 (**Figure S5D**), and had low relative expression of previously identified reactive astrocyte genes (**Figure 4D**, **Supplemental file 1**). Of these defining genes, Arhgef4 (Rho guanine nucleotide exchange factor 4) has been reported to play a role in G protein coupled receptor-mediated responses to extracellular stimuli. In addition, Arhgef4 suppresses neuronal function by sequestering PSD95, and its ablation is associated with enhancement in hippocampal-dependent spatial memory in the object location memory test (*32*). Rgs6 also encodes a protein associated with regulation of G protein signaling (*33*). SNPs of Rgs6 have been implicated in numerous psychiatric disorders (*34*) and its overexpression promotes inflammation and oxidative stress in spinal cord injury (*35*). Lastly, Enox1, involved in plasma membrane electron transport pathways, is associated with memory deficits in patients with psychiatric conditions (*36*). Thus, existing evidence supports that astrocyte cluster A7 contains a population of disease-enhancing astrocytes that promote inflammation and neuronal dysfunction in Arctic mice, and that are completely suppressed by inhibition of C5aR1 signaling.

Astrocyte clusters were analyzed for genes associated with pan-reactive, A-1 neurotoxic, or A-2 astrocytes (*31*). The WT-enriched clusters A1 and A2 had very low expression of reactive genes (**Figure 4D**). In contrast, cluster A3, highly elevated in all Arctic samples, showed high expression of pan reactive and “A-2” astrocyte genes, demonstrating these as response to injury independent of C5aR1 signaling. Interestingly, pan-reactive markers Lcn2 and Cp (ceruloplasmin) were more highly expressed in Arc-PMX205 astrocytes compared to Arc-Veh, while A-1 genes H2-T23 and H2-D1 were strongly downregulated in Arc-PMX205 astrocytes. Overall, while the previously identified pan-reactive, A-1, and A-2 genes were enhanced in Arc astrocytes, PMX205 treatment substantially decreased only A1 H2-T23 and H2-D1, and muted A-2 Tm4sf1 and panreactive Serpina3n, Cxcl10 and S100a6 (**Figure 4E**). However, PMX205 had a profound effect on a neurotoxic cluster of astrocytes that arise in Arctic mice, cluster A7, as well as rescuing neuronal supportive cluster A6 and enhancing protective cluster A10, indicating that many disease-associated astrocytes are not induced when C5aR1 signaling is inhibited.

The PMX205-mediated downregulation of reactive microglia and astrocyte genes was assessed by qPCR with a younger (7 mo) cohort of Arc mice with qPCR (**Figure S6**). While PMX205 treatment did not significantly affect hippocampal levels of microglial marker Cst7 or Itgax, starting treatment at 4 mo of age for 12 weeks, reduced levels of Tyrobp (p < 0.05), Inpp5d (p < 0.0001) Lcn2 (p < 0.01), Tnf (p < 0.05) and Ccl4 (p < 0.001) compared to non-treated Arc mice were seen. Delaying the initiation of treatment and for a shorter period of time (6 weeks) did not result in a significant reduction of inflammatory gene expression, with the exception of the astrocyte gene LCN2, although 2 weeks of treatment show rapid reduction of Tyrobp and Ccl4 (p < 0.05). While reduced expression of microglial genes with PMX205 treatment was generally consistent with single cell RNA-seq data, expression of reactive astrocyte marker *Lcn2* decreased in the younger cohort treated from 4.5 to 7 months, but not in the Arc mice treated with PMX205 from 7.5 to 10 months. These data suggest that while not all “injury” responses to pathology are blocked by suppression of C5aR1 signaling, excessive glial activation (reflected here by *Tyrobp* and *Ccl4*) can be targeted even in later stages of the disease by C5aR1 inhibition.

### PMX205 treatment promotes neurotrophic and neuroprotective signaling pathways in Arctic mice

Given that C5aR1 inhibition has selective suppressive effects on glial gene expression in the hippocampus of Arctic mice, CellChat was used to infer the probability and strength of intercellular communications between different cell populations based on genotype and treatment groups (**Figure S7A-D**). OPCs were the greatest contributor to cell-cell signaling regardless of genotype or treatment (**Figure S7E-H**), highlighting their role monitoring and responding to their environment (*37*). Not surprisingly, the number of intercellular signals and the strength of those signals was greater in the Arctic mouse relative to the wildtype, and both the number and strength of interactions was reduced by inhibition of C5aR1 in the Arctic mouse (**Figure S7I-J**).

To further interrogate altered intercellular signaling in Arctic mice with and without PMX205 treatment, we used CellChat to plot predicted relative information flow of significantly altered pathways in Arc vs WT and in Arc vs Arc-PMX205 cells (**Figure 5**). Of the pathways that were significantly altered between WT and Arc cells, several were unique to the Arctic hippocampus and not found in WT cells, including Tenascin, ANGPT, FGF, MPZ, PROS, CDH5, and AGRN (**Figure 5A**). The greater number of pathways observed in Arc compared to WT cells is reflected by higher relative number of interactions (**Figure S7I**). Other pathways were relatively over-represented in Arctic cells compared to WT. These include CLDN, FN1, VTN, SEMA4, Collagen, CDH, EGF, Laminin, MAG, and PTN. Conversely, the number of pathways that were suppressed in Arc cells compared to WT were very few (CSF, PECAM1, and NRG). This is in accordance with CellChat data reporting an increase in the number of interactions between different cell populations in Arc compared to WT (**Figure S7J**). Treatment with PMX205 suppressed several of these pathways in Arctic mice (**Figure 5B**). Of the pathways that were unique to Arc cells, Tenascin, ANGPT, PROS, and AGRN were suppressed by C5aR1 antagonism, while FGF, MPZ, and CDH5 were further enhanced by treatment. We identified three additional pathways that were unique to Arc-PMX205 cells, which were SEMA7, TGFß, and BMP.

**Fig.5:**
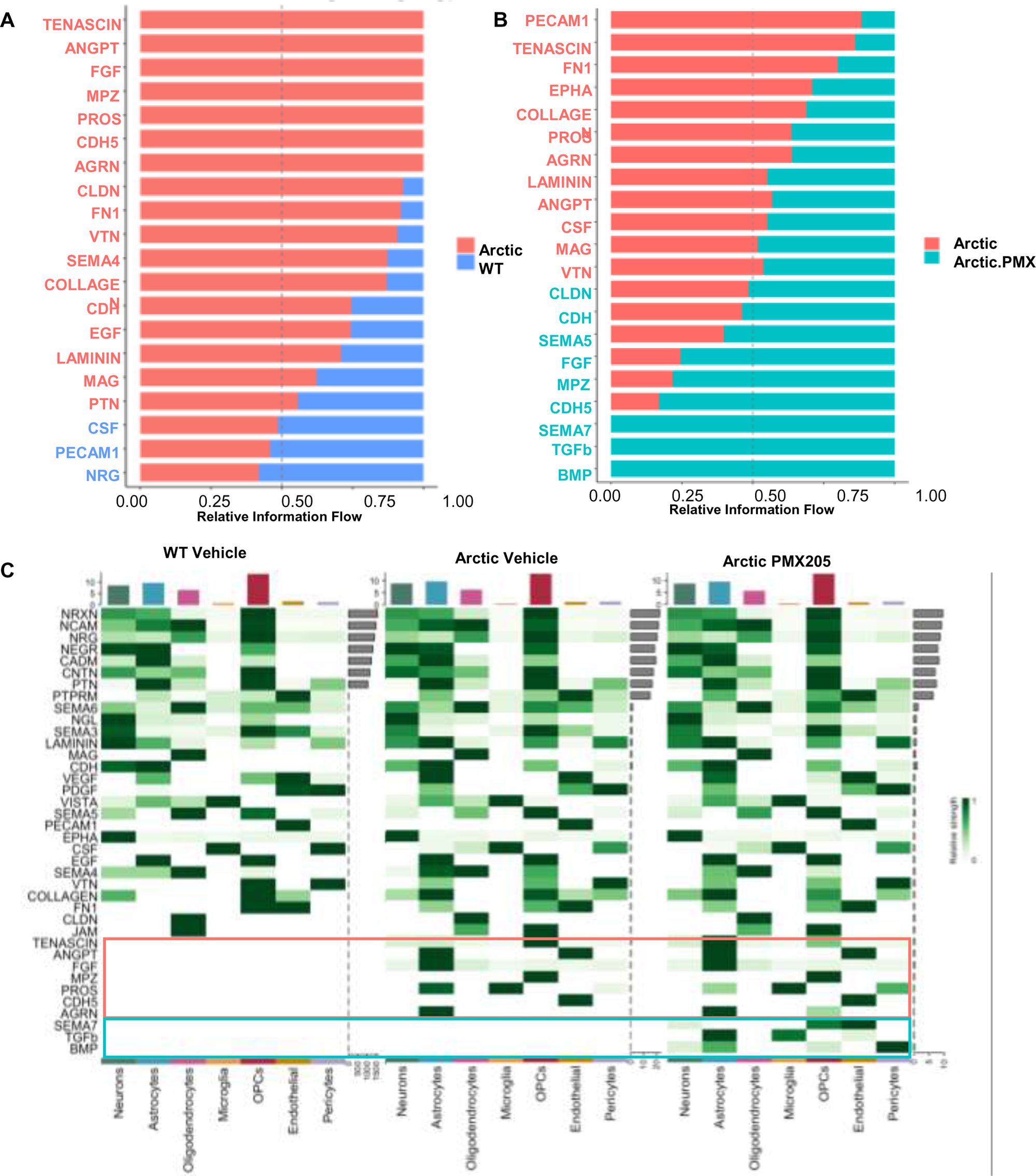
Relative information flow of pathways significantly enhanced or suppressed in Arctic mice and altered by PMX205 treatment. (**A**) Relative information flow of pathways altered between WT-veh (blue) and Arc-veh (salmon). (**B**) Relative information flow of pathways altered between Arc-veh (salmon) and Arc-PMX (teal). (**C**) Cellular senders of pathways altered in WT-veh (left), Arc-veh (middle), and Arc-PMX (right).

Interestingly, the majority of signaling that was enhanced in Arctic cells compared to WT cells derived from astrocytes (**Figure 5C**). These included ANGPT, FGF, PROS, and AGRN (**Figure 5C**, orange boxes). PROS signaling was also observed from microglia (**Figure 5C**). Similarly, PMX205 treatment enhanced signaling pathways derived from astrocytes (**Figure 5C**, blue boxes), including BMP and TGFß. Signaling was also enhanced from OPCs and endothelial cells. Many pathways were preserved between Arc-Veh and Arc-PMX205 cells. These pathways had a higher signaling probability and play important roles in cell adhesion and are known to bind to the extracellular matrix (NCAM, NRXN, NRG, CADM, NEGR, CNTN, and PTN) to allow for synaptic stabilization and organization (NRXN, NRG, and PTN) (**Figure S8**). The specific ligand-receptor interactions between different cell types were also defined to determine the senders and receivers of specific signaling pathways (**Figure S9**). For example, we identified enhanced FGF1 signaling from Arc-PMX astrocytes to neurons, microglia, oligodendrocytes, OPCs, endothelial cells, pericytes, and other astrocytes (**Figure S9**).

### PMX205 induces dynamic changes in glial markers in the Arctic hippocampus

To determine how pharmacological inhibition of C5aR1 signaling influenced plaque and glial pathology, we used immunohistochemistry to assess plaque, microglia, and astrocyte markers in the hippocampus of PMX205 treated vs untreated Arctic mice. Confocal imaging of the CA1 region of the hippocampus did not show differences in the pan-microglial marker Iba1. In both Arc-Veh and Arc-PMX, Iba1+ microglial staining was abundant. However, immunostaining for markers of reactive microglia such as CD11b and CD11c, which make up part of the integrin receptors CR3 and CR4, respectively, revealed significant increases (63% and 45%, respectively) in the CA1 region in response to PMX205 treatment (**Figure 6A-D**). It appears that C5aR1 inhibition in the Arc mouse suppresses upregulation of DAM1 genes while allowing an increase of DAM2-type proteins, at least some of which may have beneficial roles (*38*).

**Fig.6:**
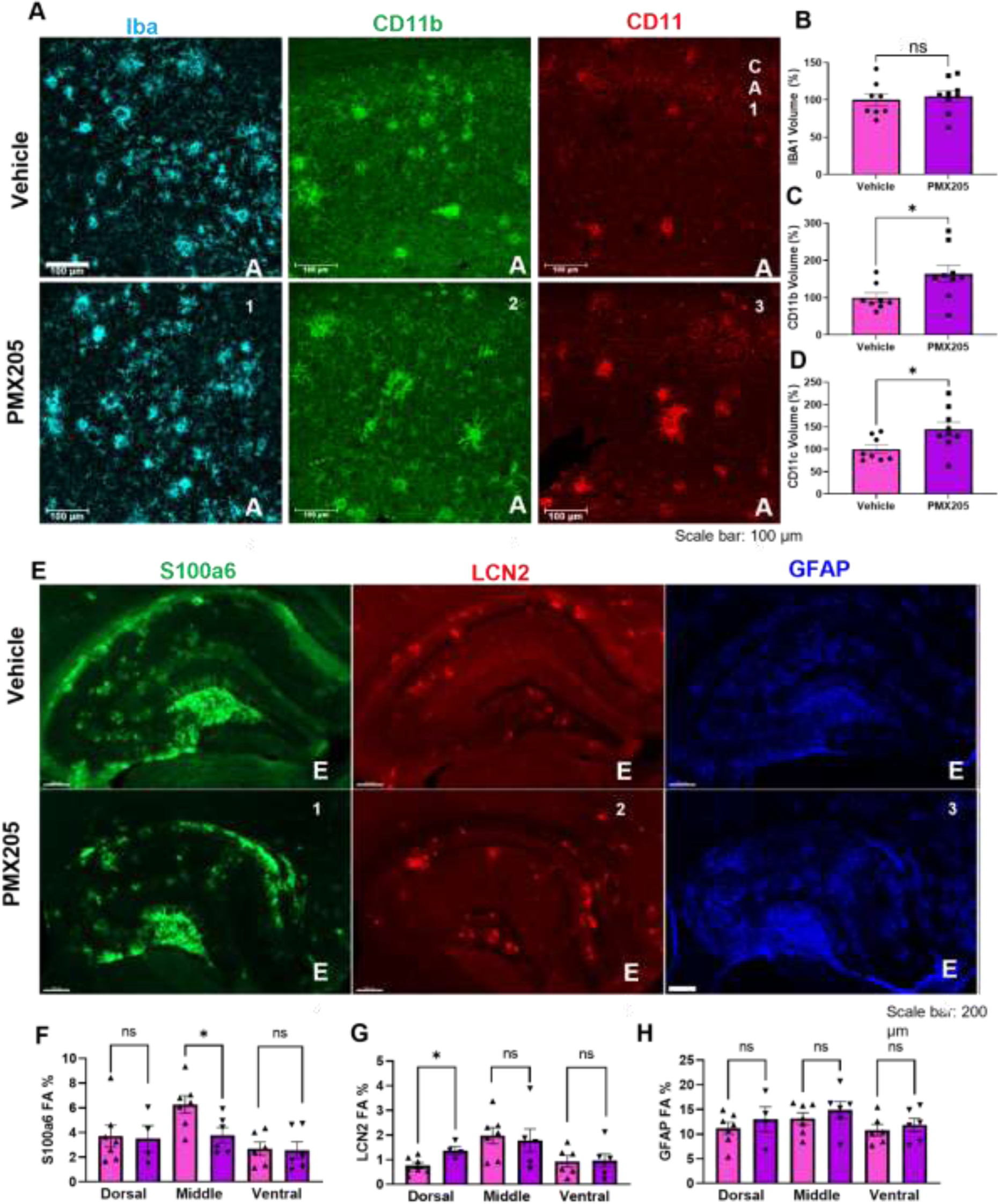
PMX205 induces dynamic changes in hippocampal microglia and astrocyte protein markers: (**A**) Representative images of hippocampal region CA1 stained for Iba1 (**A_1_, A_4_**), CD11b (**A_2_, A_5_**) and CD11c (**A_3_, A_6_**) in Arc-veh (top panel) and Arc-PMX (bottom panel) 20X magnification, scale bar 100 μm. (**B-D**) Quantification of staining volume normalized to Arc-veh levels for Iba1 (**B**), CD11b (**C**), and CD11c (**D**). (**E**) Representative images of dorsal hippocampus stained for S100a6 (**E_1_, E_4_**), LCN2 (**E_2_, E_5_**), and GFAP (**E_3_, E_6_**) in Arc-veh (top panel) and Arc-PMX (bottom panel) 10X magnification, scale bar 200 μm. (**F-H**) Quantification of percent field area of S100a6 (**F**), LCN2 (**G**), and GFAP (**H**) was done separately in sections of the dorsal, middle, and ventral hippocampus. Data shown as mean ± SEM. * *p* < 0.05, t-test (**B-D**) and One-way ANOVA with Tukey’s *post hoc* (**F-H**). N = 4-7 mice/genotype/treatment.

To assess changes in astrocytes, we stained dorsal, middle, and ventral hippocampal slices from Arc mice with antibodies reactive with S100a6, LCN2, and GFAP (**Figure 6E-H**). While PMX205 treatment had no effect on hippocampal GFAP levels, it did result in a 40% decrease in S100a6 field area localized to the middle hippocampal section (consistent with RNA-seq data and relevant as S100a6 levels are elevated in inflammatory diseases including AD (*39*)). Interestingly, a 44% increase in LCN2 field area specific to the dorsal hippocampus was detected, again compatible with RNA-seq data (**Figure 4E**). Mice that were treated from 7.5 to 10 months did not have a reduction in AmyloGlo staining (**Figure S10**).

### Inhibition of C5a-C5aR1 signaling protects short-term spatial memory in female Arctic mice

Finally, to determine if the altered hippocampal signaling induced by C5aR1 inhibition is associated with hippocampal-dependent memory in Arc mice, we used the Y maze spatial reference test (**Figure 7**) to test male and female mice. Female Arc mice spent significantly less time exploring the novel, previously blocked arm of the Y maze compared to WT mice (19% vs 29%, *p* = 0.025). However, this deficit was eliminated in female Arc mice treated with PMX205, who spent 47% of the trial in the novel arm and significantly outperformed Arc-Veh mice (*p* = 0.0014) (**Figure 7B**). Interestingly, Y maze performance did not change in male mice depending on treatment [*F* (1, 37) = 0.09, *p* = 0.52] or genotype [*F* (1, 37) = 0.42, *p* = 0.87], with most male mice exploring the novel arm 40-44% of the time allotted (**Figure 7C**), suggesting that disease progression and the onset of memory deficits is slower in male Arctic mice. The accelerated deficit in hippocampal-dependent 336 memory in female Arc mice (prevented by C5aR1 antagonism) compared to male mice reflects the overexpression of DAM1 genes in microglia derived from female Arc mice compared to male Arc mice (**Figure S3**). Finally, plasma NfL levels were measured before and after PMX205 treatment in mice treated at early time points (4.5-6.5 months to 7 months of age) and later time points (7.5 to 10 months of age). In the younger cohort, Arc-Veh mice had higher levels of NfL compared to WT mice at 7 months of age (p < 0.0001) and compared to their pre-treatment levels (at 4.5 months). Treatment with PMX205 for 2 (p < 0.05), 6 (p < 0.01), or 12 (p = 0.13) weeks prevented this time-dependent increase (**Figure 8A**). In the older cohort, Arc mice at 7.5 months of age had higher levels of plasma NfL compared to age-matched WT mice (**Figure 8B**). At the end of treatment, Arc mice still had higher levels of plasma NfL than the WT, but there was no difference between Veh- and PMX205-treated Arc mice (**Figure 8B**). These findings suggest that inhibition of C5aR1 may limit NfL levels in plasma in Arc mice in early stages of plaque accumulation, but not at later ages when plaque deposition and neurotoxicity is more extensive.

**Fig.7:**
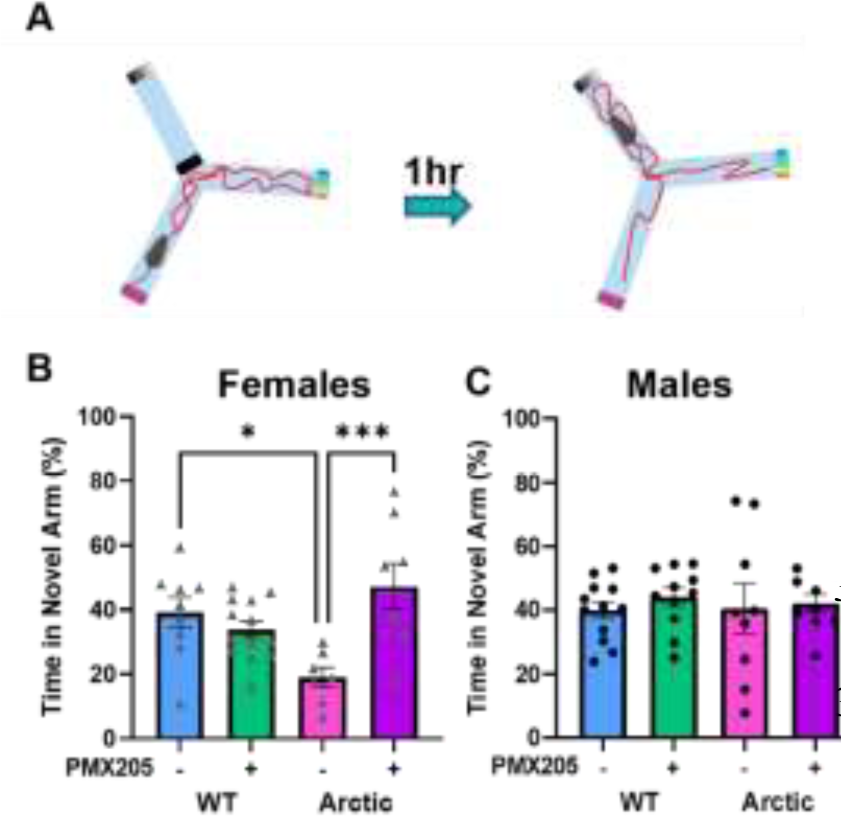
PMX205 protects against spatial memory deficits in Arctic females. (**A**) Overview of Y maze experiment. (**B-C**) Time spent in the novel arm during the test trial in females (**B**) and males (**C**). Data shown as mean ± SEM. * *p* < 0.05; *** *p* < 0.001, Two-way ANOVA with Tukey’s *post hoc.* N = 6-13 mice/sex/genotype/treatment.

**Fig.8:**
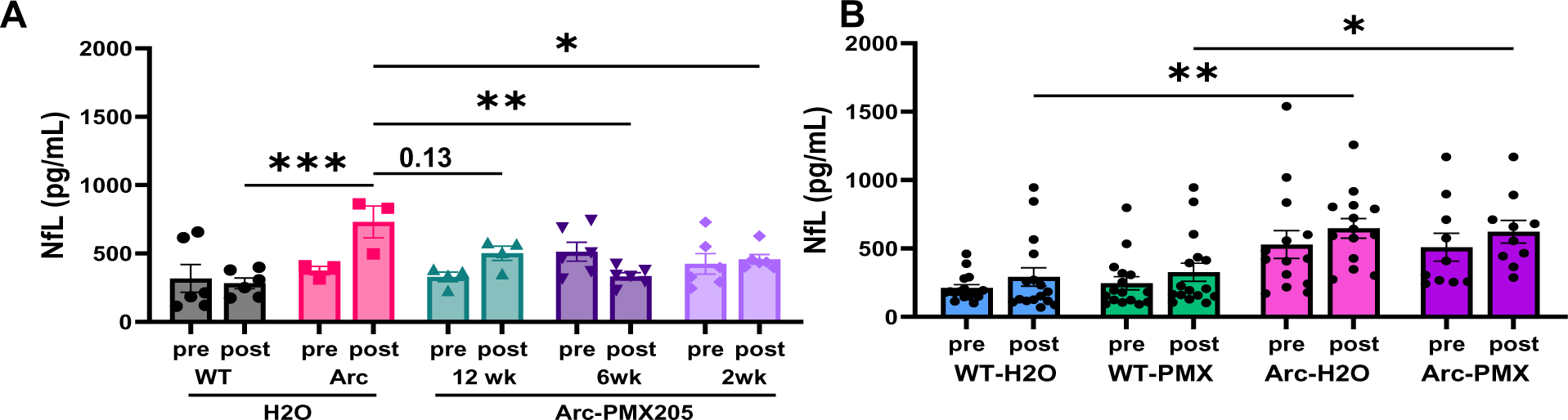
Early treatment with PMX205 suppresses neurofilament light levels in plasma of Arc mice. (**A**) Plasma NfL levels in 4.5 mo WT and Arc prior to treatment and in 7 mo WT and Arctic mice after PMX205 treatment for 2, 6, or 12 weeks compared to WT and Arctic age-matched controls. (**B**) Plasma NfL levels of WT and Arctic mice before and after PMX205 or Vehicle (Veh) treatment from 7.5 to 10 months. Data shown as mean ± SEM. * p < 0.05; ** p < 0.01; *** p < 0.001, Repeated measures Two-way ANOVA with Dunnet’s post hoc. N = 3-6 mice/treatment (**A**) and Two-way ANOVA with Sidak’s post hoc, N = 10-16 mice/genotype/treatment (**B**). Males and females were included in both studies with no apparent sex differences evident in any genotype/treatment group.

## DISCUSSION

Complement activation contributes to proper synaptic connectivity and plasticity during development as well as in adults (*40, 41*). However, overactivation or dysregulation of complement can result in excessive synaptic pruning, neuroinflammation and cell death. Activation of the classical complement pathway can be triggered by interaction of C1q in the C1 complex by fibrillar Aβ plaques, neurofibrillary tangles as well as the exposure of damage associated molecular patterns on neuronal cell surfaces (*42–45*), all of which can lead to downstream production of C5a and C3a. C5aR1 and C3aR are prominently induced in microglia in response to injury (as reviewed in (*46*) and confirmed in our model (*19*)). In brain, if complement is activated and C5 is present locally to enable production of C5a, C5aR1 signaling can initiate pro-inflammatory patterns of gene expression. In the periphery, C5a-C5aR1 signaling is linked to pro-inflammatory responses such as chemotaxis, degranulation, and cytokine production in multiple disease models. C5aR1 also works synergistically with toll-like receptors to enhance inflammatory responses (*47–49*). Here, we demonstrate that pharmacologic inhibition of C5a-C5aR1 signaling improved cognitive outcomes in the aggressive Arctic mouse model of AD, and, while increasing some protein markers associated with microglial reactivity (CD11b/CD11c), decreased transcription of disease-enhancing microglial genes and altered astrocyte polarization.

The findings presented here expand our previous investigations of the effects of PMX205 treatment in the Tg2576 (*20, 21*) and 3xTg (*21*) models to the very aggressive Arctic model. While inhibition of C5aR1 signaling suppressed several parameters of glial activation in all models, in the slower progressing Tg2576 and 3xTg models, amyloid plaques accumulation was decreased by 50%, while that was not observed in the Arctic mice. However, functional protection was seen in both the Tg2576 and Arctic models (behavior was not assessed in the 3xTg), consistent with the hypothesis that inhibition of C5aR1 dampens the detrimental inflammatory response including that to Aβ plaques, rendering them less toxic despite not limiting plaque volume. Genetic ablation of C5aR1 in Arctic mice also showed rescue of behavior, maintenance of neuronal integrity and either a decrease or delay in expression of AD related, DAM, and reactive astrocyte-associated genes again without substantially reducing the accumulation of amyloid plaques (*17, 19*). In these distinct studies, we used bulk RNA sequencing of dissected hippocampus and cortex (*19*), RNA sequencing of microglia isolated from brain (*17*), single cell RNA sequencing of microglia isolated from of hippocampus or cortex (*20*), and here both single cell RNA-seq of microglia and single nuclei RNA-seq from hippocampus to probe with increasing depth the molecular basis for the neuroprotective effects that result from the inhibition of C5a-C5aR1 signaling. Single nucleus RNA-seq clearly demonstrated upregulation of cell specific transcription for individual complement proteins in the Arctic mouse, consistent with the patterns detected by bulk sequencing, but demonstrating expression in specific clusters of cell type subsets that are induced by pathology. C1q was predominantly induced by microglia, C3 in astrocytes, and C4b (the mouse C4 gene) in both astrocytes and oligodendrocytes, supporting earlier reports of these individual components (*31, 50–53*). C5 (Hc) was detected in our samples at very low levels and in a small numbers of neurons and astrocytes, but it has been detected in low amounts by others in astrocytes and neurons (*53, 54*). In contrast, almost all clusters of neurons expressed the complement control proteins Csmd1 and Csmd2, suggesting a cell autonomous regulation of complement activation by neurons. Both Csmd1 and Csmd2 have been associated with multiple disorders including those associated with cognition (*55–57*). Importantly, specific microglial and astrocyte clusters were identified as displaying age/pathology dependent functional consequences of the expression of the APP transgene that were C5aR1-independent and those that were dependent on C5aR1 signaling. Microglia clusters M4 and M9 were C5aR1 dependent and expressed genes that modulated synapse pruning, microglial activation, cognition and cell death, whereas clusters M3 and M8 which had high expression of genes previously characterized as DAM-2 genes were induced in the Arctic without C5aR1 input, and may have repair or restorative functions, which one would optimally not want to suppress. In addition, M2 which contributes to organization of synapses is suppressed in C5aR1-sufficient Arctic mice but is rescued by C5aR1 inhibition. This is in line with previous data showing that C5aR1 inhibition in Tg2576 mice reduces expression of synaptic pruning genes and attenuates engulfment of synaptic puncta by microglia (*20*) and with a wide range of human and mouse studies demonstrating that some microglial responses are attempts to protect from injury (reviewed in (*58, 59*)). Similarly, A3 astrocytes were induced in response to the APP transgene regardless of C5aR1 function, while A6, associated with axon guidance and synapse transmission, was suppressed in Arctic, but rescued with inhibition of C5aR1 signaling and A7 was seen only in Arctic without PMX205 treatment. Disease specific changes in astrocyte gene expression have also been reported in SN RNA-seq data from human AD samples (*60*).

While C5aR1 expression is dominant in activated microglia and peripheral myeloid cells, CellChat analysis of hippocampal SN RNA data revealed that inhibition of C5a-C5aR1 signaling in Arctic mice affected several signaling pathways associated with not only microglia, but also astrocytes and oligodendrocytes, consistent with a complex cellular phase of AD (*61*), that may be blunted by this targeted approach in at least some subtypes of AD (*62*). CellChat revealed that PMX205 treatment in Arc mice promoted signaling of fibroblast growth factors (FGFs) from astrocytes. FGFs comprise a large family of polypeptides that are essential for cell development, maintenance, and repair (*63*). FGFs promote astrocyte differentiation and proliferation (*64*). In models of spinal cord injury, FGF2 is secreted by astrocytes to signal back to other astrocytes for regulating astrogliosis and promoting structural changes in glia. Importantly, it was shown that FGF signaling is important for the return to homeostasis of reactive astrocytes following acute CNS injury. FGF activation suppresses excessive autophagy and promotes clearance of misfolded protein aggregates via PI3K/Akt/mTOR signaling (*65*). Elevation of FGF in astrocytes after spinal cord injury also promotes survival of OPCs (*64*). FGF2 has been shown to be downregulated in the prefrontal cortex of AD patients (*66*). In the APP23 mouse model of AD, overexpression of FGF2 is neuroprotective, restoring hippocampal-dependent spatial memory and reducing levels of Aβ and the APP cleaving enzyme (BACE) (*66*). Attenuation of cognitive decline by FGF may involve stimulation of hippocampal neurogenesis (*67*). Interestingly, FGF stimulation increased the number of hippocampal astrocytes without affecting levels of pan-reactive microglial marker Iba1 (*66*), suggesting that in this context, astrocytes may contribute to neuronal health and survival. However, *in vitro* studies show that glutamate- or oligomeric Aβ-induced toxicity promotes FGF2 secretion by neurons, and this event promotes microgliosis and phagocytosis of neuronal debris via ERK signaling (*68*). These findings suggest that the neuroprotective effects of FGF2 may be context- and injury-dependent and different sources of FGF2 (i.e., exogenous vs endogenous) may activate either astrocytes or microglia to promote cell growth or debris clearance, respectively. Furthermore, complement activation can directly stimulate FGF2 release from retinal pigment epithelial cell cultures via formation of MAC (*69*). By targeting complement activation at C5aR1, MAC formation in the Arc mouse model may contribute directly to FGF signaling. Thus, it is possible that PMX205 treatment in Arctic mice promotes FGF signaling from astrocytes, which then downregulates inflammatory signaling from astrocytes and promotes proliferation of neuroprotective astrocytes. These findings support the consideration of FGF pathway stimulation as a therapeutic approach to combat neuronal death and promote cell growth in AD, perhaps in conjunction with complement C5aR1 inhibition.

TGFβ2 signaling from astrocytes to microglia and endothelial cells was also enhanced in PMX205 treated Arctic mice. TGFβ is a super family of proteins that includes TGFβ, BMPs, and activins. TGFβ binding to its receptor activates SMAD3, which promotes amyloid-β uptake and iNOS production by microglia and astrocytes, respectively (*70*). TGFβ signaling is important for microglial development and promotes expression of quiescent microglial genes including *P2ry12, Fcrls*, and *Sall1* (*70*). Mice lacking TGFβ1 in the CNS fail to develop ramified microglia and instead only have microglia with amoeboid or “activated” morphology (*70*). Furthermore, microglia act via TGFβ to promote hippocampal neurogenesis in inflammatory or neurodegenerative contexts (*71*). BMPs are also members of the TFGβ superfamily of signaling ligands that play a role in CNS development (*72*). BMPs promote neurogenesis and astrogliogenesis and help maintain adult neural stem cells in the subgranular zone (SGZ) of the hippocampus (*72*). BMP5 is highly expressed by pericytes and its signaling promotes astrocyte survival (*73*). Cell-cell communication from pericytes to astrocytes was increased in Arc mice with PMX205 treatment relative to vehicle-treated cells as is the BMP pathway, supporting the possibility that with C5aR1 inhibition, pericytes send trophic or survival factors, such as BMP5, to astrocytes. The increased pericyte to astrocyte signaling was not as robust in WT mice suggesting that the increased signaling is a response to inflammatory conditions. We found BMP5 signaling to be present in pericytes isolated from Arc-PMX hippocampi, with signaling directed towards astrocytes, neurons, and OPCs. Notably, this pathway was absent in Arc-Veh cells. Finally, evidence from preclinical models of ischemic stroke suggest that reactive astrocytes upregulate expression of tenascin to inhibit proliferation, possibly as a self-limiting mechanism (*74*). The shift of tenascin signaling from OPCs to astrocytes in the Arc-PMX205 cells, in addition to FGF signaling, suggests that inhibition of C5aR1 promotes a return to homeostatic state in astrocytes exposed to amyloid pathology.

The complete genetic knock out of C5aR1 resulted in protection from loss of spatial memory (*17*). The results presented here show that pharmacologic inhibition of C5aR1 in adults with substantial amyloid plaque load can also protect against hippocampal-dependent spatial memory deficits, highlighting its potential as a therapeutic target to slow or ameliorate cognitive decline in adults. While male Arc mice did not exhibit a deficit at the age tested (10 mo), female Arc mice showed a prominent deficit in short-term spatial memory in the Y maze that was attenuated with PMX205 treatment. Sex differences have not been reported in the long-term cognitive performance of Arctic mice (*17, 19*), however, in the present study female Arctic mice had more robust inflammatory microglial gene expression compared to males (with the same plaque accumulation), suggesting that enhanced inflammation may contribute to spatial memory decline in Arctic mice. AD is more prevalent in women than men, and women with AD also show a steeper decline in memory compared to men with AD. Male and female rodents also have differences in hippocampal structure and neurogenesis (*75*) and have been reported to use distinct strategies for spatial navigation (*75, 76*), and thus, it is possible that this short-term memory test was more sensitive to detect sex differences. Lastly, modulating the onset of treatment may have greater potential for reducing neuronal injury and cognitive decline. While suppression of plasma NfL, a biomarker of neurodegeneration, was noted at 7 mo of age after 2-12 weeks of PMX205 treatment, unexpectedly, plasma NfL at 10 months of age were not impacted by C5aR1 inhibition. Thus, C5aR1 inhibition via PMX205 may be most effective if initiated at earlier stages of the disease for maximum effect.

C5aR1 has become a target of interest for several neuroinflammatory conditions. Clinical and preclinical studies reported increased complement proteins in the brain and CSF in amyotrophic lateral sclerosis, stroke, epilepsy, traumatic brain injury, multiple sclerosis, and Alzheimer’s disease (reviewed in (*4, 77*)). For example, elevated C5a is associated with white matter lesions in multiple sclerosis (*78*), severity of traumatic brain injury (*79*), and increased Aβ-induced neuronal injury (*80*). Interestingly, evidence supports that the less understood C5a receptor 2 (C5aR2) prevents lesion formation and promotes recovery after spinal cord injury (*28*). Such studies suggest that targeting the pro-inflammatory receptor C5aR1 would be more beneficial than targeting the ligand C5a, which may interact with C5aR2 to promote healing. It is critical to note that complement is an important protective system against infection, clearing dead cells and debris (*81*), and thus complete inhibition of the system, especially C1q through C3, would have significant immunocompromising effects. The 2021 FDA approval of the small molecule C5aR1 antagonist, avacopan (Tavneos) for treatment of the peripheral disorder, antineutrophil cytoplasmic antibody-associated vasculitis, as well as earlier small studies of PMX53 in humans, suggests a lack of toxicity when blocking this receptor in humans. Whether the newly approved avacopan or PMX205 will be effective in substantially slowing the progression of cognitive decline in Alzheimer’s disease or other neurodegenerative disorders awaits human clinical trials. In summary, in this study, treatment with PMX205 when plaque accumulation is robust was sufficient to attenuate the inflammatory response to the plaques and preserve short-term spatial memory. Combined with data from several animal models and positive effects in humans, inhibition of C5aR1 reduces inflammatory responses while preserving neuroprotective functions of complement activation. Thus, targeting C5aR1 has high potential for treatment of neurodegenerative diseases, and thus is a promising candidate for clinical trials in disorders such as AD.

## MATERIALS AND METHODS

### Animals

The Institutional Animal Care and Use Committee of University of California at Irvine approved all animal procedures and experiments were performed according to the NIH Guide for the Care and Use of laboratory animals. Mice were single-housed in ambient temperature and given access to food and water *ad libitum.* The Arctic48 mouse model of AD (hereafter referred to as Arc), which carries the human APP transgene with three mutations – the Indiana (V717F), the Swedish (K670N + M671L), and the Arctic (E22G), was generated on a C57BL6/J background and originally provided by Dr. Lennart Mucke (Gladstone Institute, San Francisco, CA). Mice were crossed with C57BL6/J wild type (WT) mice to create Arc+/- and Arc-/- (WT) mice. This hemizygous mouse model produces fibrillar plaques as early as 2 to 4 months of age (*19, 82*).

### PMX205 Treatment

C5aR1 antagonist PMX205 (Mimotopes, Victoria, Australia) was diluted in MilliQ H_2_O to a final concentration of 20 µg/ml. To fully solubilize PMX205 from powder form, we dissolved 5 mg in 1mL of MilliQ H_2_O, then diluted to the final concentration. Drinking bottles were filled with 100 mL of PMX205 or vehicle (MilliQ H_2_O) and were weighed and refilled weekly. Mice were single housed throughout treatment in order to calculate drug consumption for each individual mouse. Mice were also weighed weekly to detect potential toxic effects of PMX205 treatment.

In a smaller pilot experiment, Arc mice were treated with PMX205 via drinking water for 2, 6, or 12 weeks beginning at 6.5, 5.5 or 4.5 months of age, respectively to detect biomarker changes. A cohort of age-matched WT and Arc mice were untreated and used as controls. For all cohorts, plasma samples were collected prior to treatment by submandibular puncture under anesthesia and at perfusion via cardiac puncture. Brains were perfused and collected for IHC, and hippocampi were dissected for RNA-seq and qPCR assays.

### Y maze spatial reference test

Mice were subjected to behavioral testing. Female mice were tested before male mice, and equipment was thoroughly cleaned or changed between sex groups to eliminate odor cues. The Y maze test for spatial memory reference was adapted from (*83*) to measure short-term hippocampal-dependent spatial memory. Briefly, mice were placed into an arm of a three-armed maze, with one of the arms blocked off and given 5 minutes to explore, after which mice were placed back in their home cages. After a 1-hour intertrial interval, the block from the third arm was removed, and mice were again given 5 minutes to explore freely. Time spent in each arm was tracked with Noldus EthoVision. We compared total time spent in the novel arm between groups and preference for the novel arm over the two familiar arms within groups with Two-way ANOVA and Tukey’s post hoc test. This test took place in normal (200 Lux) lighting. The ends of each arm had distinct visual patterns so that mice could distinguish between a novel and familiar arm. The mazes were cleaned with 70% ethanol and dried thoroughly after each trial.

### Neurofilament light assay

Blood was collected before treatment via submandibular puncture and before perfusion via cardiac puncture for plasma isolation. Blood was immediately mixed with 0.2 M EDTA to a final concentration of 10 mM and centrifuged at 5,000rpm (Eppendorf Centrifuge) for 10 minutes at 4°C. Supernatant (plasma) was collected and stored in −80°C until use. To measure plasma neurofilament light (NfL) levels, the MesoScale Diagnostics NfL kit (F217X-3) was used following manufacturer’s instructions.

### Microglia isolation and fixation

Half hippocampi from 2-3 mice of the same sex, genotype and treatment were pooled for processing (3 samples per sex/genotype/treatment). To isolate microglia from hippocampi, we followed modified protocols from Miltenyi Biotech as previously described (*20, 84*). Cells were incubated in CD11b beads, and labeled microglia were collected in LS columns and then eluted from the columns. Microglia were centrifuged at 500 xg for 10 min at 4°C, resuspended in 375 µL of cell buffer from the kit with 0.5% BSA, and counted before continuing to the cell fixation protocol provided by Parse Biosciences (V1.3.0). After incubation in cell permeabilization solution, 2 mL of cell neutralization buffer were added to each sample and they were centrifuged at 500 xg for 10 minutes at 4°C. Pellets were resuspended in cell buffer with DMSO (1:20 dilution), counted again, aliquoted, and stored at −80°C until ready for barcoding.

### Nucleus isolation and fixation

Nucleus isolation was done with Nuclei Extraction Buffer (Miltenyi Biotech 130-128-024) following manufacturer’s instructions. Briefly, perfused half hippocampi (3 per sex/genotype/treatment) were collected and frozen at −80° until use. Frozen hippocampi were placed directly into C-tubes with 2 mL lysis buffer (Nuclei Extraction Buffer with 0.2 U/µL RNase inhibitor) and processed in the Miltenyi gentleMACS^TM^ Octo Dissociator. Nuclei were passed through a 70 µm SmartStrainer. An additional 2 mL of lysis buffer was added to the C-tubes and passed through the strainer to capture more nuclei. Samples were then centrifuged at 350 xg at 4°C for 5 minutes. Nuclei were resuspended in 1.5 mL of resuspension buffer (PBS with 0.1% BSA and 0.2 U/µL RNase inhibitor), passed through a 30 µm filter, and counted before proceeding to nuclei fixation protocol (Parse Biosciences V1.3.0). 4 million nuclei were collected from the stock of each sample preparation, centrifuged, and resuspended in 750 µL nuclei buffer with 0.75% BSA (Parse Biosciences). Samples were passed through a 40 µm filter, incubated in nuclei fixation solution for 10 minutes on ice, followed by incubation in nuclei permeabilization solution. 4 mL of nuclei neutralization buffer were added to each sample and they were centrifuged at 500 xg for 10 minutes at 4°C. Pellets were resuspended in nuclei buffer with DMSO (1:20), frozen and stored at −80°C.

### Barcoding and library preparation for single cell and single nuclei RNAseq

The Evercode Whole Transcriptome (WT) kit (Parse Biosciences V1.3.0) was used to prepare single nucleus and single cell microglia libraries for RNA-seq. Forty-eight samples (24 single cell and 24 single nucleus) diluted to 525 cells or nuclei/µL were each loaded onto a separate well in a 96-well plate provided by Parse Biosciences for initial barcoding. Following reverse transcription and barcoding, all SC and SN samples were pooled and incubated in ligation mix for a second round of barcoding. The pooled cells were then redistributed to a 96-well plate. This step was repeated once more for a total of 3 barcoding cycles. The cell/nuclei mix was then pooled again and counted to calculate final concentration. After cell counting, 7 separate aliquots of 12,500 cells or nuclei per sublibrary were processed in parallel, for a combined 87,500 cells/nuclei total. For each sublibrary, cells/nuclei were lyzed, barcoded cDNA was amplified, fragmented, and a unique sublibrary index was added prior to sequencing. The seven sublibraries were sequenced on a single NextSeq2000 (Illumina) sequencing run for an average of 10902reads/cell. Resulting Fastq files and data matrices were deposited in GEO with the accession ID GSE240950.

### RNA-seq processing and data analysis

Fastq files were processed using the Parse short-read split-pipe pipeline as previously described (*85*). Cells and nuclei were filtered using different metrics from the matrices generated. Cells were filtered for more than 300 UMIs, < 10% mitochondrial reads, < 50,000 counts and > 300 genes. Nuclei were filtered for more than 300 UMIs, < 1% mitochondrial reads, < 50,000 counts and > 400 genes. Then, the cells and nuclei were integrated with Seurat. In downstream analyses, normalization and clustering were calculated using Seurat SCTransform and Louvain algorithms, respectively (*86*). Intercellular communication between clusters of cells was inferred using CellChat (*87*).

### Quantitative PCR

Frozen hippocampi from 7 mo WT (n = 6) and Arc H2O controls (n = 4), and Arc mice treated for 2 (n = 6), 6 (n = 6), and 12 (n = 5) weeks with PMX205 were pulverized using mortar and pestle for RNA quantification of target genes as previously described (*17*). The following FAM dye and MGB quencher TaqMan probes were used (ThermoFisher): Mm00438349_m1 (Cst7), Mm00498708_g1 (Itgax), Mm01324470_m1 (Lcn2), Mm00443260_g1 (Tnf), Mm0043111_m1 (Ccl4), Mm00449152_m1 (Tyrobp), and Mm00494987_m1 (Inpp5d). All probes were multiplexed with VIC dye and MGB quencher probe Mm01545399_m1 (Hprt) as an internal control for each well. cDNA from each hippocampus was tested in triplicate. Within each well, relative expression to Hprt was calculated by subtracting the VIC cycle threshold (Ct) value from FAM Ct value. ΔCt values within triplicates were averaged then exponentially transformed and multiplied by 1000 (2^-ΔCt^ * 1000). P values were calculated using one-way ANOVA followed by Dunnet’s multiple comparisons test.

### Immunohistochemistry

Mice were deeply anesthetized with isoflurane, perfused transcardially and half brains dissected, fixed and 30 µm coronal sections were obtained for immunohistochemistry following previously reported protocols (*19*). Primary antibodies used were rabbit anti-Iba1 (1:1000, Wako #019-19741), rat anti-CD11b (1:1000, Biorad #MCA74G), hamster anti-CD11c (1:400, Biorad #MCA1369), goat anti-Lcn2 (1:50, R&D #AF1857), chicken anti-GFAP (1:1000, Abcam #ab4674), and rabbit anti-S100a6 (1:500, Novus #NBP1-89388). Alexa Fluor secondary antibodies, diluted 1:500 in blocking solution (*19, 20*), included goat anti-Armenian hamster alexa 568 (Abcam, # ab175716), goat anti-rat alexa 488 (Invitrogen #A-11006), goat anti-rabbit alexa 555 (Invitrogen #A-11070), goat anti-rabbit alexa 647 (Invitrogen #A-21244), donkey anti-goat alexa 555 (Invitrogen #A-21432), donkey anti-rabbit alexa 488 (Invitrogen #A-21206), and donkey anti-chicken alexa 647 (Jackson Immuno Research #803605155). To visualize plaques tissues were incubated for 10 min with Amylo-Glo (1:100 in PBS, Biosensis #TR-300-AG). Low magnification images (10X) were acquired using ZEISS Axio Scan.Z1 Digital Slide Scanner. Higher magnification z-stacks were acquired at 20X magnification with Leica SP8 confocal microscope. The percent volume of markers for microglia, astrocytes, and plaques were quantified using the Surfaces feature of Imarisx64 (version 9.5.0). Quantitative comparisons between groups were carried out on comparable sections of each animal processed at the same time with the same batches of solutions.

### Statistical Analysis

Power analysis was performed *a priori* to determine sample size for behavioral assays using G*Power. Statistical analyses were performed with GraphPad Prism (V 9.5.0). Treatment X genotype effects were analyzed using two-way ANOVA with Tukey’s *post hoc* test. Pre- and post-treatment effects were compared with repeated measures ANOVA with Sidak’s *post hoc* test.

## List of Supplementary Materials

Fig. S1 - S10

Table S1

## Acknowledgments

This study was made possible in part through access to the Optical Biology Core Facility of the Developmental Biology Center, a shared resource supported by the Cancer Center Support Grant (CA-62203) and Center for Complex Biological Systems Support Grant (GM-076516) at the University of California, Irvine.

## Funding

National Institutes of Health grant R01 AG060148 (AJT and AM)

Alzheimer’s Association Research Fellowship AARFD-20-677771 (NDS)

Larry L. Hillblom postdoctoral fellowship #2021-A-020-FEL (AGA)

T32 AG00096 (TJP)

Edythe M. Laudati Memorial Fund.

## Author contributions

Conceptualization: NDS, AJT, AM

Methodology: NDS, HYL, KC, SC, AMA, TJP, AGA, AM, AJT

Investigation: NDS, HYL, KC, SC, AMA, TJP, AGA

Visualization: NDS, HYL, KC, SC

Funding acquisition: NDS, AJT, AM, AGA, TJP

Project administration: NDS, AM, AJT, SC, HYL Supervision: AJT, AM

Writing – original draft: NDS, AJT

Writing – review & editing: NDS, HYL, KC, SC, AMA, TJP, AGA, AM, AJT

## Data and materials availability

The code used in this study will be made available upon request. All transcriptomic data are available in the supplementary materials. Fastq files and data matrices were deposited in GEO with the accession ID GSE240950.

