## Supplemental Figures 1-10 for "C5aR1 antagonism suppresses inflammatory glial gene expression and alters cellular signaling in an aggressive Alzheimer’s model"

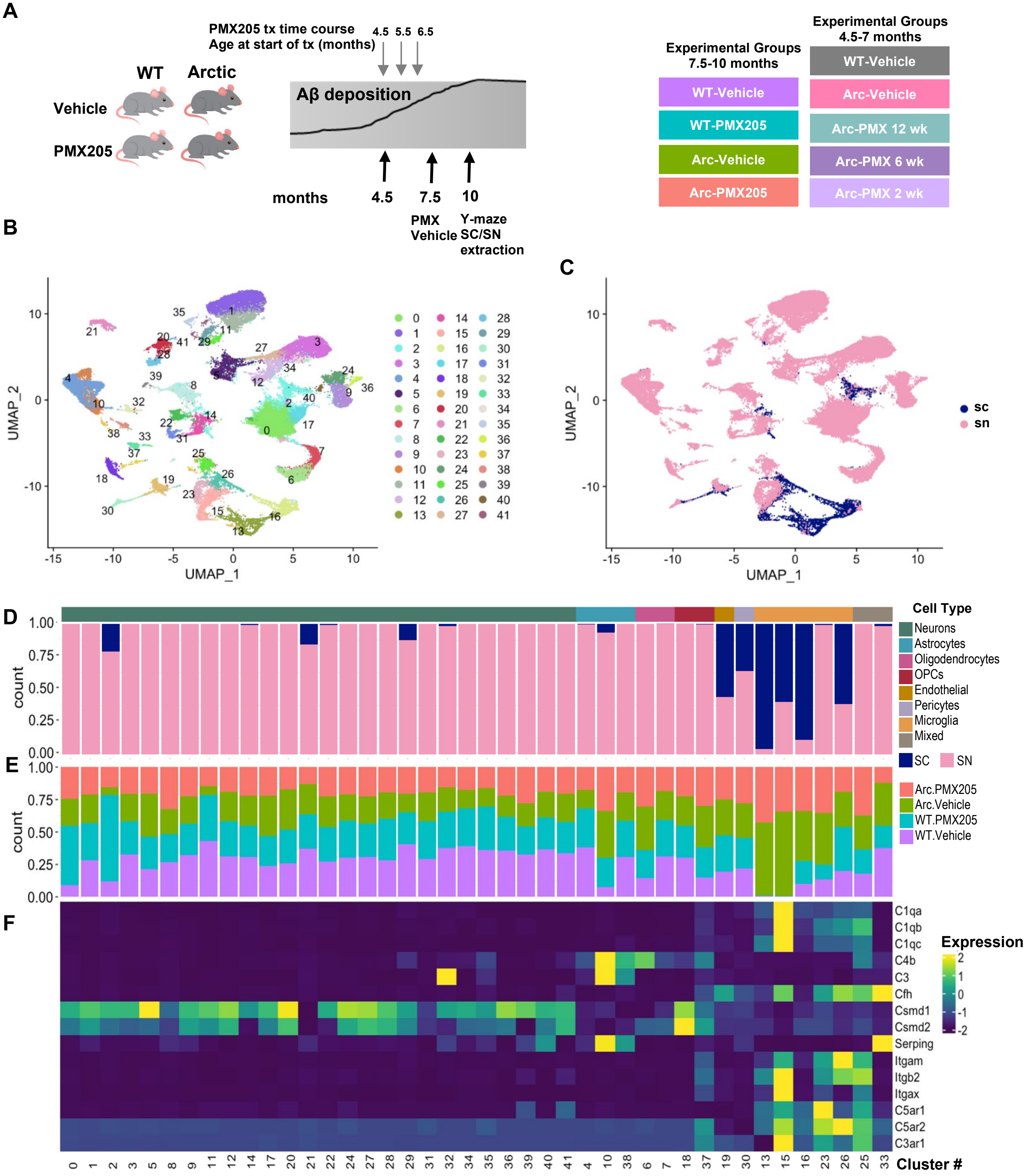

**Fig.1: Clustering of transcriptome from single nucleus and single cells in wild type and Arctic mice.** (**A**) Experimental timeline for PMX205 treatments (**B**) Seurat cluster identification of combined cells/nuclei transcriptome. (**C**) Seurat cluster identification of cells derived from single cells (SC) or single nucleus (SN) sequencing. (**D**) Proportion of cells in each cluster originating from SC or SN transcriptome. (**E**) Proportion of cells in each cluster by treatment/genotype. (**F**) Expression of complement pathway components, receptors or regulators by cell type.

(**Figure 1F**). The astrocyte cluster that was enriched in Arc samples expressed high levels of C3, C4b, and Serping1.

Of the 42 Seurat clusters, 26 were identified as neurons, 5 as microglia, and 3 as astrocytes (**Figure 1B, 1D**). Among the single nucleus samples, neurons made up the majority of cell types sequenced from each of the treatment groups. We also identified 2 clusters of oligodendrocytes and 2 of oligodendrocyte progenitor cells (OPCs), as well as single clusters of endothelial cells, pericytes, and two "mixed" clusters that are likely a mixture of astrocytes and microglia (**Figure 1D and 2A, 2B**).

Because we used a treatment to modulate complement activation-mediated signaling, we further explored the relative complement gene expression in each cell cluster and genotype/treatment group **(Figure S2A).** Neuronal clusters expressed relatively high levels of complement regulatory genes Csmd1 and Csmd2 (irrespective of genotype), and astrocytes had high transcript levels of C4b, C3 and Serping1 (C1 Inhibitor) expressed predominantly in Arc mice (**Figure 2C**). Microglial clusters expressed C1qa, C1qb, C1qc, Cfh (Factor H), Itgam (CD11b), Itgb2 (CD18), Itgax (CD11c), C5ar1, C5ar2, and C3ar1 (**Figure 2C and 1F**). When only looking at SN samples, neuronal, endothelial, and OPC expression of complement genes was unchanged,

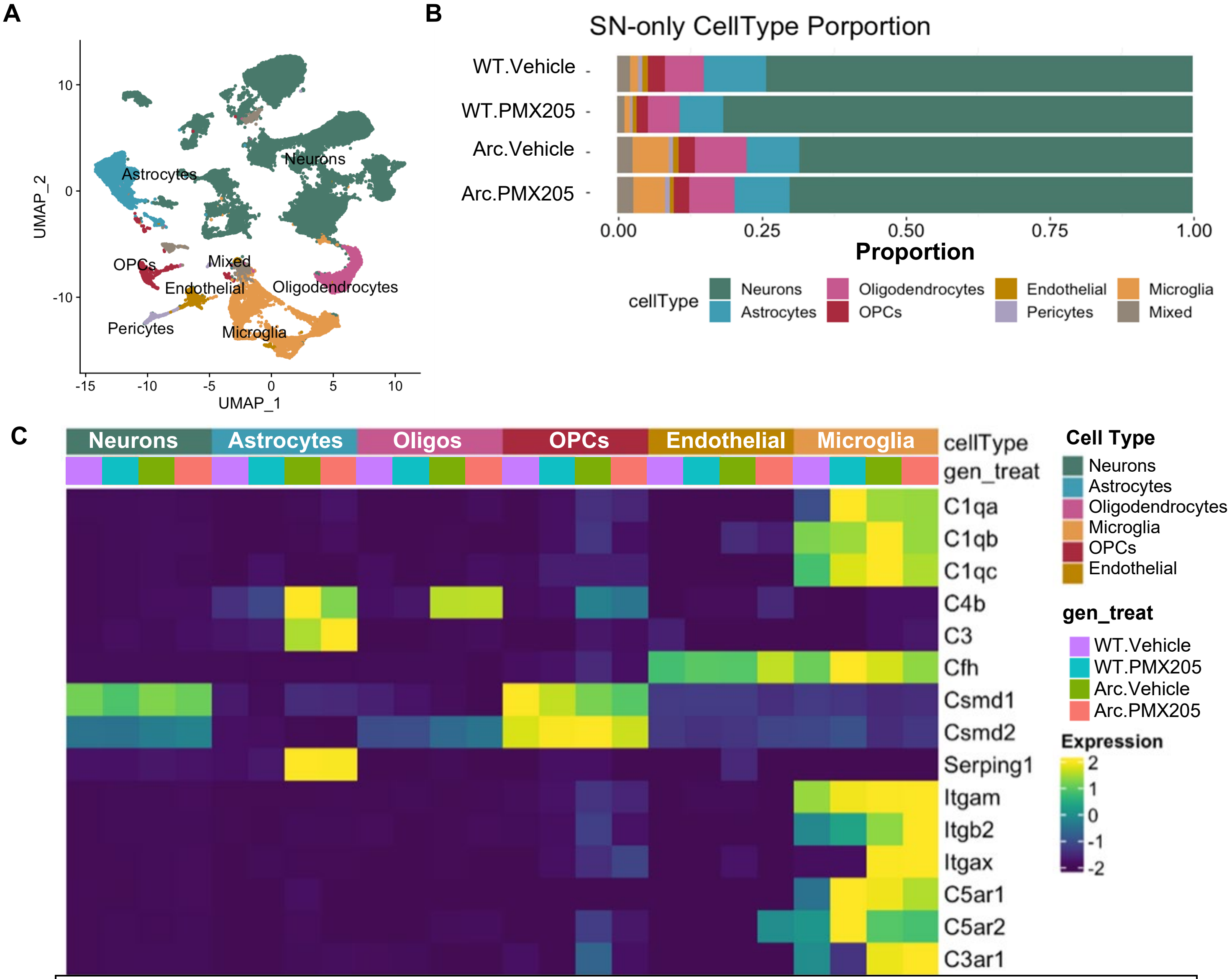


**Fig.2: Single cell and single nucleus RNA-Seq clustered by cell type reveals cell-specific complement gene expression.** Microglia or nuclei were isolated from hippocampi, fixed, and sequenced. (**A**) U-Map of all cell types with all treatment groups included (counts per cell type were: Neurons 37512, Astrocytes 4612, Oligodendrocytes 3478, Microglia 4891, OPCs 1259, Endothelial 898, Mixed 1017, Pericytes 490). (**B**) Proportion of isolated cell types in the single nucleus RNA-seq by genotype and treatment group. (**C**) Differential cell type-specific expression of complement pathway components and regulators of the complement system derived from single nucleus RNA-seq.

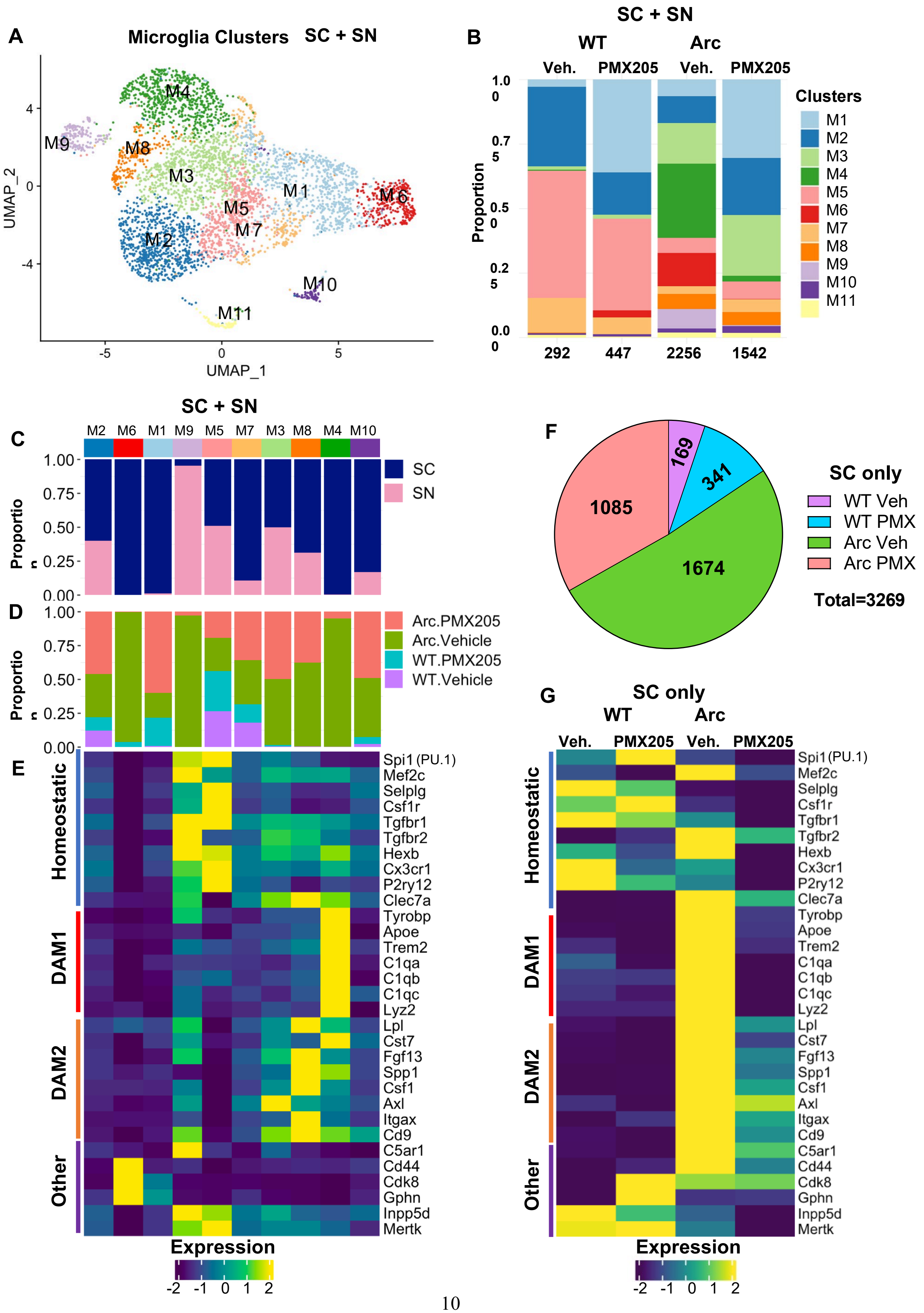
A
Microglia Clusters SC + SN
M4
M9
M8
M3
M1
M6
M5
M7
M2
M10
M11
UMAP_2
UMAP_1
B
SC + SN
WT
Arc
Veh.
PMX205
Veh.
PMX205
Proportion
1.00
0.75
0.50
0.25
0.00
292
447
2256
1542
Clusters
M1
M2
M3
M4
M5
M6
M7
M8
M9
M10
M11
C
SC + SN
M2 M6 M1 M9 M5 M7 M3 M8 M4 M10
SC
SN
Proportion
D
Arc.PMX205
Arc.Vehicle
WT.PMX205
WT.Vehicle
Proportion
E
Homeostatic
DAM1
DAM2
Other
Spi1 (PU.1)
Mef2c
Selplg
Csf1r
Tgfbr1
Tgfbr2
Hexb
Cx3cr1
P2ry12
Clec7a
Tyrobp
Apoe
Trem2
C1qa
C1qb
C1qc
Lyz2
Lpl
Cst7
Fgf13
Spp1
Csf1
Axl
Itgax
Cd9
C5ar1
Cd44
Cdk8
Gphn
Inpp5d
Mertk
Expression
F
SC only
169
341
1085
1674
WT Veh
WT PMX
Arc Veh
Arc PMX
Total=3269
G
SC only
WT
Arc
Veh.
PMX205
Veh.
PMX205
Homeostatic
DAM1
DAM2
Other
Expression

**Fig.3: DAM1 gene expression is suppressed in Arctic-PMX205 hippocampal microglia.** (**A**) Cells/nuclei identified as microglia were re-clustered separately. (**B**) Proportion of microglia (SC+SN) clusters in WT-Veh, WT-PMX, Arc-Veh, and Arc-PMX samples. (**C**) Proportion of cells in each cluster originating from SC or SN transcriptome. (**D**) Proportion of cells/nuclei in each cluster by treatment/genotype. (**E**) Relative expression of genes representative of homeostatic microglia, DAM1, DAM2, or other genes of interest within the different microglial clusters. (**F**) Pie chart demonstrating proportion of SC microglia samples derived from different treatment groups. (**G**) Relative expression of homeostatic, DAM1, or DAM2 genes in different treatment groups, with SC data alone.

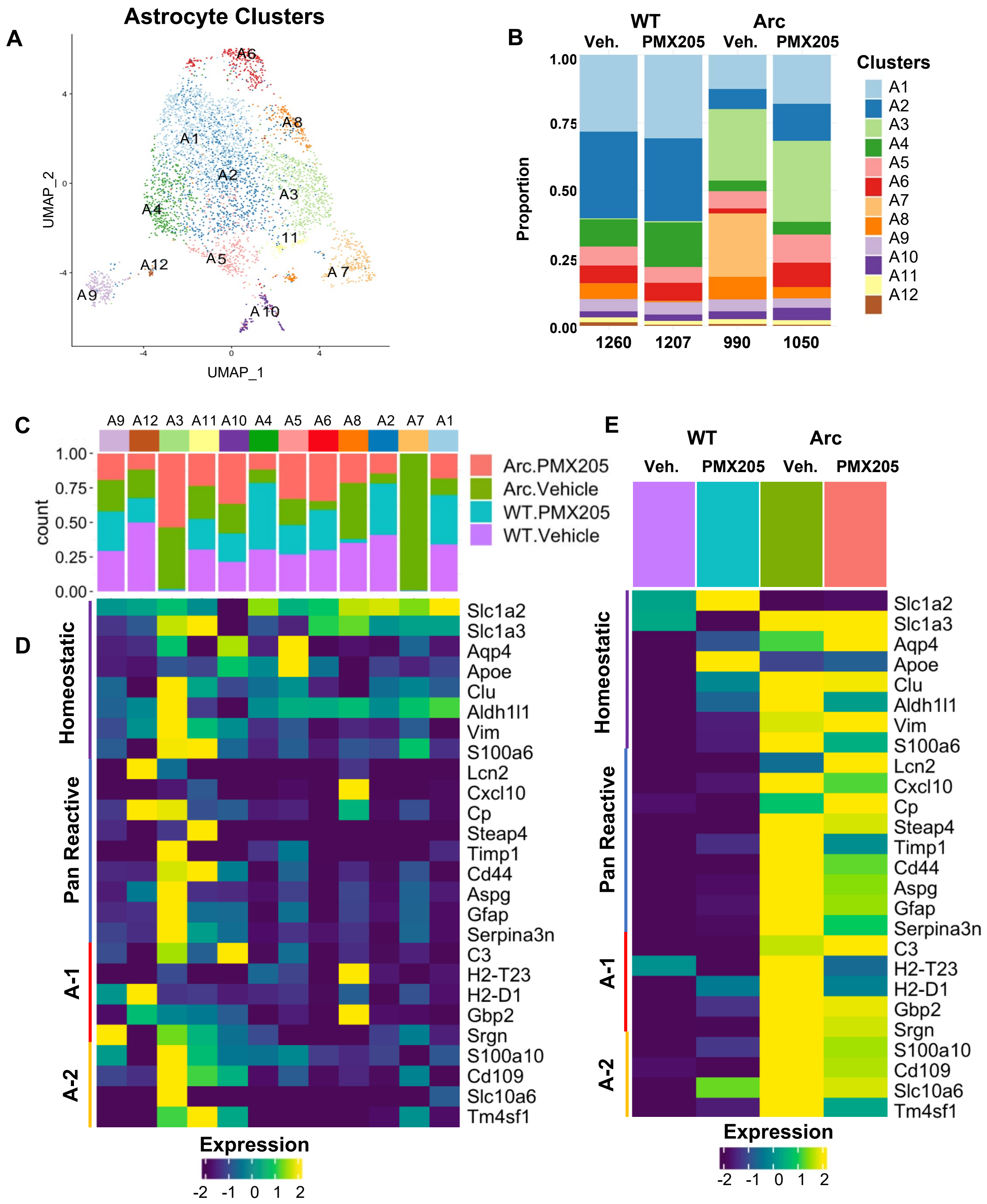


signaling is inhibited that results in support of neuronal functions in response to inflammatory stimulus.

**Fig.4: Reactive Astrocyte gene expression is largely suppressed in Arctic-PMX205 hippocampus.** (**A**) Cells identified as astrocytes were re-clustered separately. (**B**) Proportion of astrocyte populations in WT-Veh, WT-PMX, Arc-Veh, and Arc-PMX samples. (**C**) Proportion of cells in each cluster by treatment/genotype. (**D**) Relative expression of genes representative of homeostatic astrocytes, pan-reactive, A-1 neurotoxic, or A-2 neuroprotective astrocytes within the different astrocyte clusters. (**E**) Relative expression of homeostatic, pan-reactive, A-1, or A-2 genes in the different treatment groups.

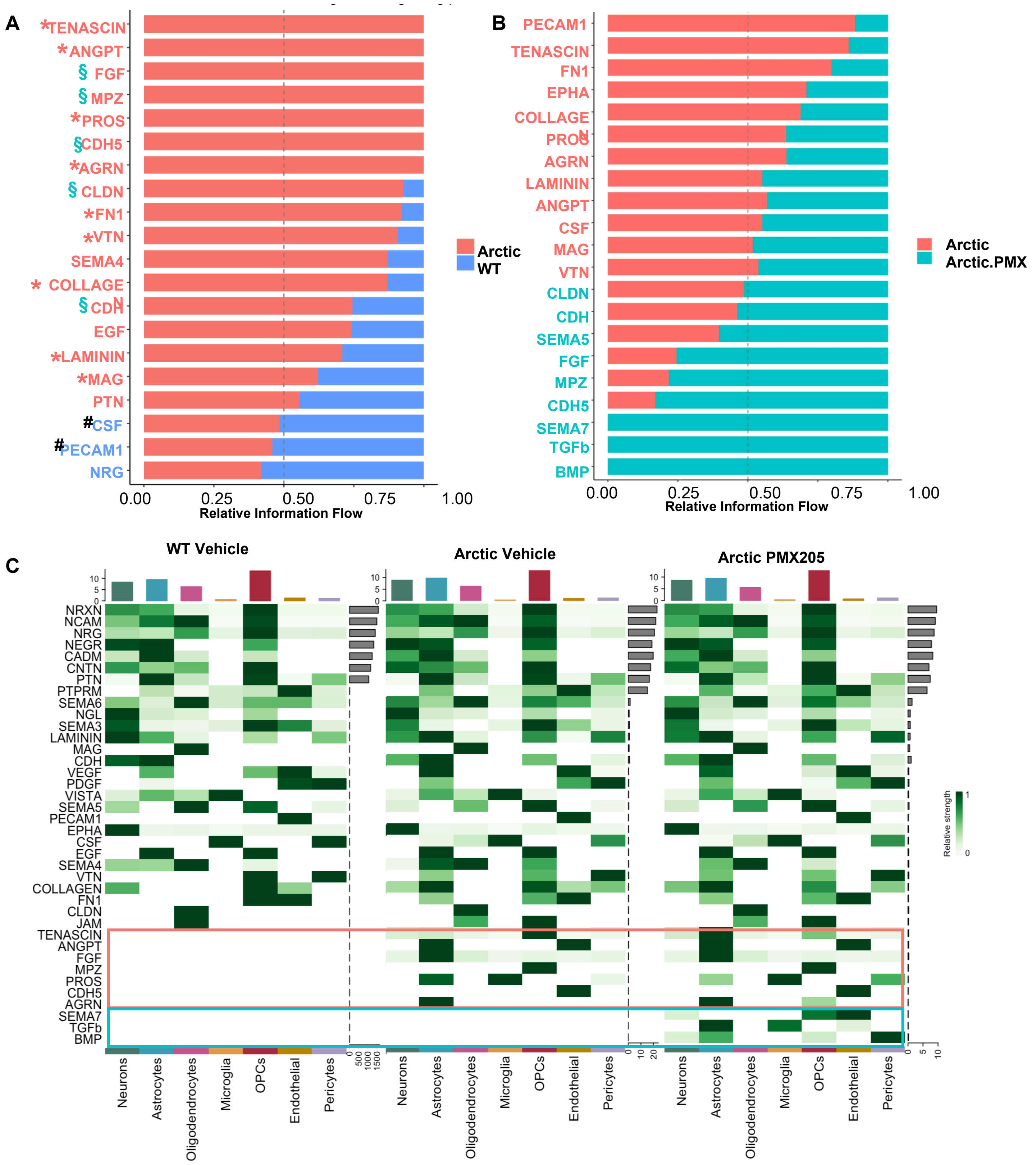

**Fig.5: Relative information flow of pathways significantly enhanced or suppressed in Arctic mice and altered by PMX205 treatment.** (**A**) Relative information flow of pathways altered between WT-veh (blue) and Arc-veh (salmon). (**B**) Relative information flow of pathways altered between Arc-veh (salmon) and Arc-PMX (teal). (**C**) Cellular senders of pathways altered in WT-veh (left), Arc-veh (middle), and Arc-PMX (right).

suppressed several of these pathways in Arctic mice (**Figure 5B**). Of the pathways that were unique to Arc cells, Tenascin, ANGPT, PROS, and AGRN were suppressed by C5aR1 antagonism, while FGF, MPZ, and CDH5 were further enhanced by treatment. We identified three additional pathways that were unique to Arc-PMX205 cells, which were SEMA7, TGFß, and BMP.

Interestingly, the majority of signaling that was enhanced in Arctic cells compared to WT cells derived from astrocytes (**Figure 5C**). These included ANGPT, FGF, PROS, and AGRN (**Figure 5C**, orange boxes). PROS signaling was also observed from microglia (**Figure 5C**). Similarly, PMX205 treatment enhanced signaling pathways derived from astrocytes (**Figure 5C**, blue boxes), including BMP and TGFß. Signaling was also enhanced from OPCs and endothelial cells. Many pathways were preserved between Arc-Veh and Arc-PMX205 cells. These pathways had a higher signaling probability and play important roles in cell adhesion and are known to bind to the extracellular matrix (NCAM, NRXN, NRG, CADM, NEGR, CNTN, and PTN) to allow for synaptic stabilization and organization (NRXN, NRG, and PTN) (**Figure S8**). The specific ligand-receptor interactions between different cell types were also defined to determine the senders and receivers of specific signaling pathways (**Figure S9**). For example, we identified enhanced FGF1 signaling from Arc-PMX astrocytes to neurons, microglia, oligodendrocytes, OPCs, endothelial cells, pericytes, and other astrocytes (**Figure S9**).

**PMX205 induces dynamic changes in glial markers in the Arctic hippocampus**

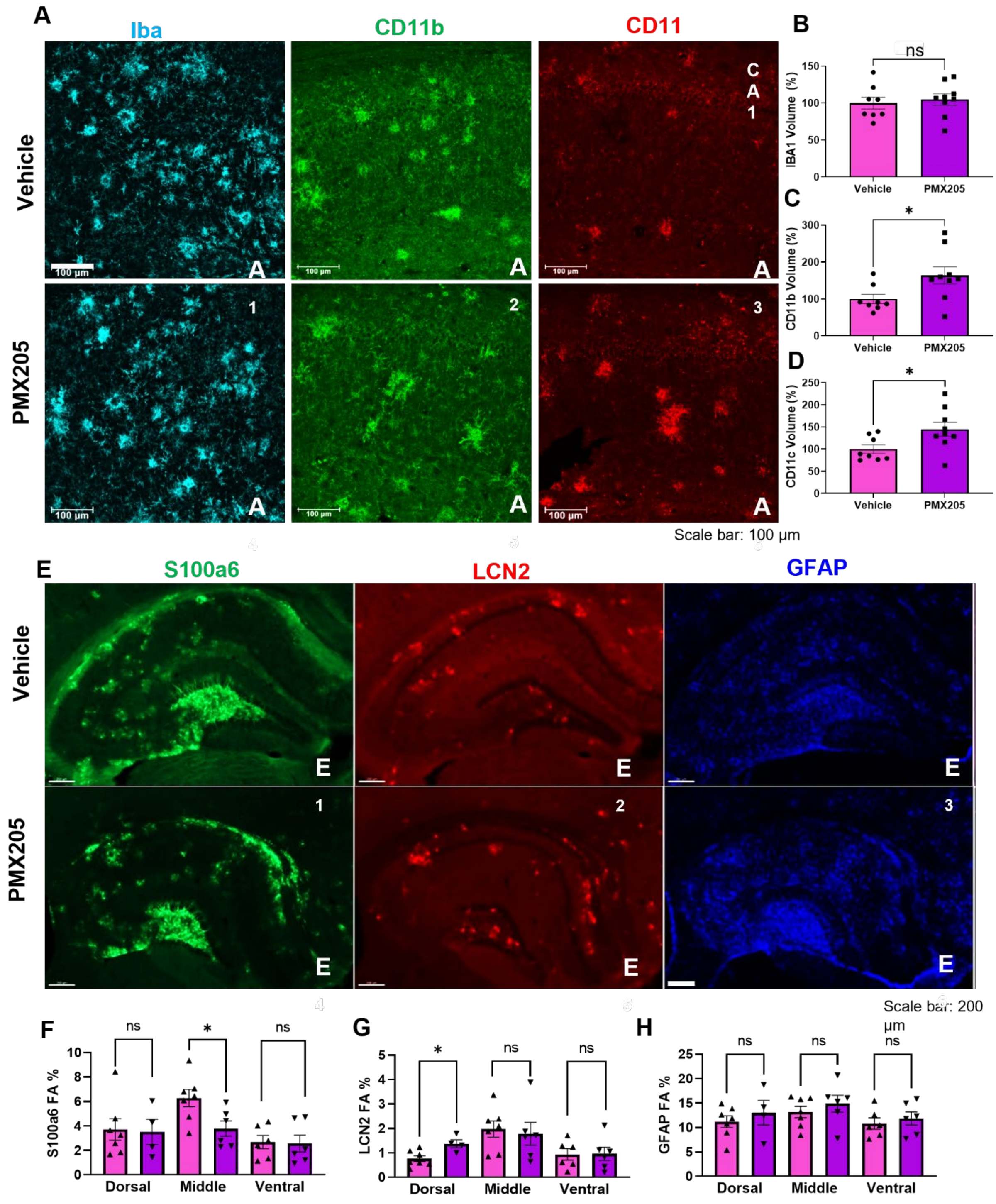


To determine how pharmacological inhibition of C5aR1 signaling influenced plaque and glial pathology, we used immunohistochemistry to assess plaque, microglia, and astrocyte markers in the hippocampus of PMX205 treated vs untreated Arctic mice. Confocal imaging of the CA1

**Fig.6: PMX205 induces dynamic changes in hippocampal microglia and astrocyte protein markers:** (**A**) Representative images of hippocampal region CA1 stained for Iba1 ($\mathbf{A_1}$, $\mathbf{A_4}$), CD11b ($\mathbf{A_2}$, $\mathbf{A_5}$) and CD11c ($\mathbf{A_3}$, $\mathbf{A_6}$) in Arc-veh (top panel) and Arc-PMX (bottom panel) 20X magnification, scale bar 100 µm. (**B-D**) Quantification of staining volume normalized to Arc-veh levels for Iba1 (**B**), CD11b (**C**), and CD11c (**D**). (**E**) Representative images of dorsal hippocampus stained for S100a6 ($\mathbf{E_1}$, $\mathbf{E_4}$), LCN2 ($\mathbf{E_2}$, $\mathbf{E_5}$), and GFAP ($\mathbf{E_3}$, $\mathbf{E_6}$) in Arc-veh (top panel) and Arc-PMX (bottom panel) 10X magnification, scale bar 200 µm. (**F-H**) Quantification of percent field area of S100a6 (**F**), LCN2 (**G**), and GFAP (**H**) was done separately in sections of the dorsal, middle, and ventral hippocampus. Data shown as mean ± SEM. * $p < 0.05$, t-test (**B-D**) and One-way ANOVA with Tukey's *post hoc* (**F-H**). N = 4-7 mice/genotype/treatment.

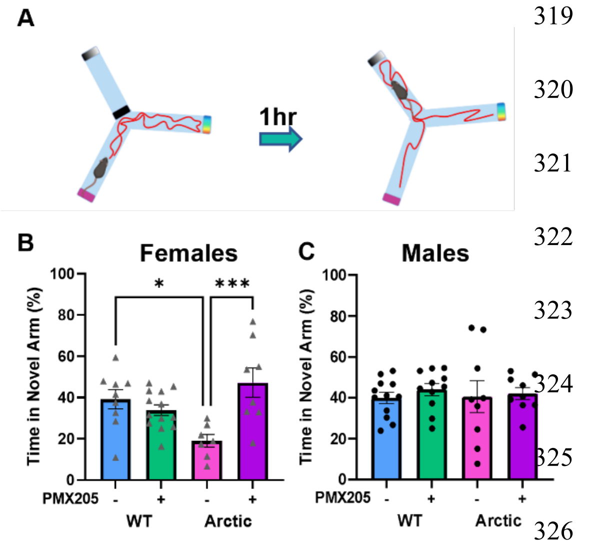


**Fig.7: PMX205 protects against spatial memory deficits in Arctic females.** (**A**) Overview of Y maze experiment. (**B-C**) Time spent in the novel arm during the test trial in females (**B**) and males (**C**). Data shown as mean ± SEM. * $p < 0.05$; *** $p < 0.001$, Two-way ANOVA with Tukey's *post hoc*. N = 6-13 mice/sex/genotype per treatment.

C5aR1 antagonism) compared to male mice reflects the overexpression of DAM1 genes in microglia derived from female Arc mice compared to male Arc mice (**Figure S3**). Finally, plasma NfL levels were measured before and after PMX205 treatment in mice treated at early time points (4.5-6.5 months to 7 months of age) and later time points (7.5 to 10 months of age). In the younger cohort, Arc-Veh mice had higher levels of NfL compared to WT mice at 7 months of age ($p < 0.0001$) and compared to their pre-treatment levels (at 4.5 months). Treatment with PMX205 for 2 ($p < 0.05$), 6 ($p < 0.01$), or 12 ($p = 0.13$) weeks prevented this time-dependent increase (**Figure 8A**). In the older cohort, Arc mice at 7.5 months of age had higher levels of plasma NfL compared

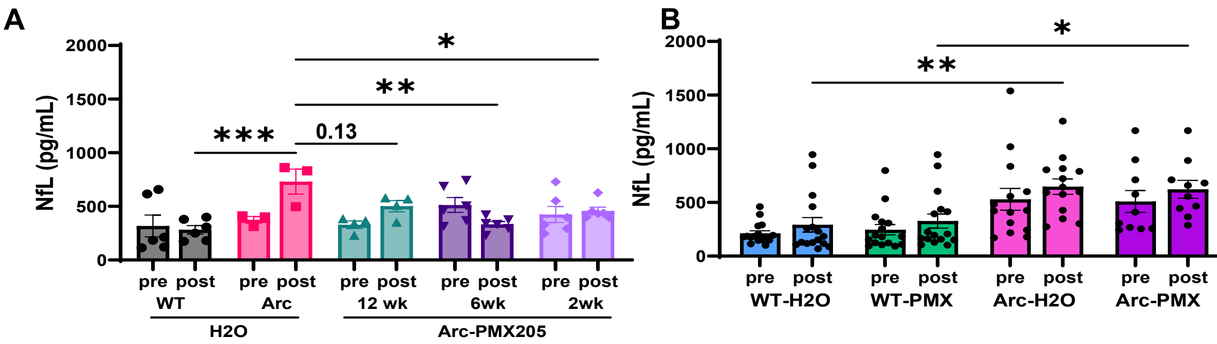


**Fig.8: Early treatment with PMX205 suppresses neurofilament light levels in plasma of Arc mice.** (**A**) Plasma NfL levels in 4.5 mo WT and Arc prior to treatment and in 7 mo WT and Arctic mice after PMX205 treatment for 2, 6, or 12 weeks compared to WT and Arctic age-matched controls. (**B**) Plasma NfL levels of WT and Arctic mice before and after PMX205 or Vehicle (Veh) treatment from 7.5 to 10 months. Data shown as mean ± SEM. * $p < 0.05$; ** $p < 0.01$; *** $p < 0.001$, Repeated measures Two-way ANOVA with Dunnet's post hoc. N = 3-6 mice/treatment (**A**) and Two-way ANOVA with Sidak's post hoc, N = 10-16 mice/genotype/treatment (**B**). Males and females were included in both studies with no apparent sex differences evident in any genotype/treatment group.

Project administration: NDS, AM, AJT, SC, HYL

Supervision: AJT, AM

Writing – original draft: NDS, AJT

Writing – review & editing: NDS, HYL, KC, SC, AMA, TJP, AGA, AM, AJT

**Competing interests:** Authors declare that they have no competing interests.

**Fig.8: Early treatment with PMX205 suppresses neurofilament light levels in plasma of Arc mice.** (**A**) Plasma NfL levels in 4.5 mo WT and Arc prior to treatment and in 7 mo WT and Arctic mice after PMX205 treatment for 2, 6, or 12 weeks compared to WT and Arctic age-matched controls. (**B**) Plasma NfL levels of WT and Arctic mice before and after PMX205 or Vehicle (Veh) treatment from 7.5 to 10 months. Data shown as mean ± SEM. * p < 0.05; ** p < 0.01; *** p < 0.001, Repeated measures Two-way ANOVA with Dunnet's post hoc. N = 3-6 mice/treatment (**A**) and Two-way ANOVA with Sidak's post hoc, N = 10-16 mice/genotype/treatment (**B**). Males and females were included in both studies with no apparent sex differences evident in any genotype/treatment group.

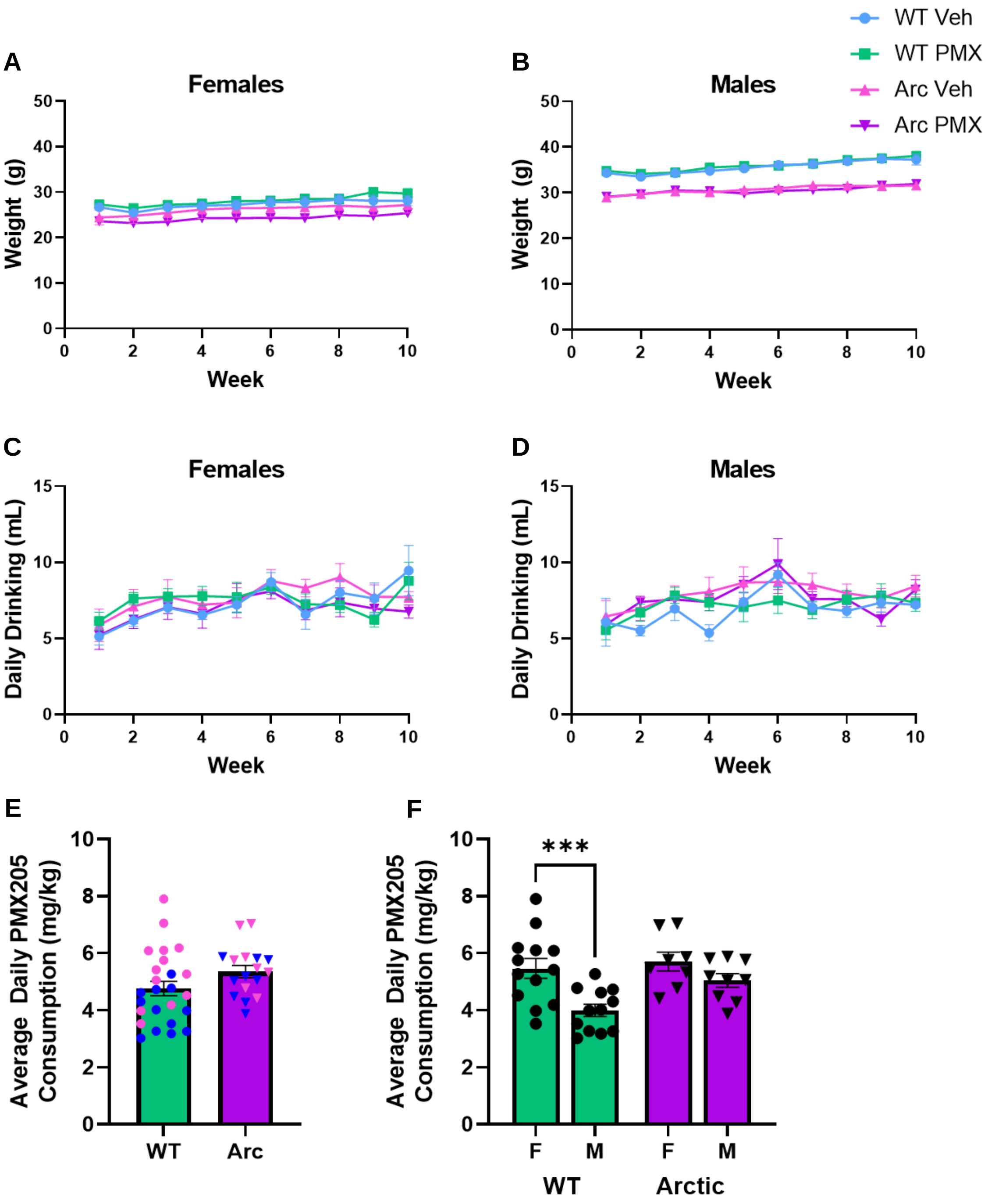


**Figure S1: PMX205 consumption did not have toxic physiological effects.** (**A-B**) Weight log (weekly) of female (**A**) and male (**B**) mice through10 weeks of treatment with or without PMX205 (**C-D**) Average volume of water consumed per week throughout treatment. (**E**) PMX205 dose was calculated based on volume consumed by each animal (females shown in pink points, males in blue). (**F**) Comparison of average daily PMX205 dose between males and females. WT males had a 26.8% smaller dose than females. Although there was no difference in volume consumed, the relative dose in males was smaller due to higher body weight. Data shown as mean ± SEM. *** $p$ <0.001, Two-way ANOVA with Sidak's *post hoc* test. N= 8-13 mice/sex/genotype/treatment

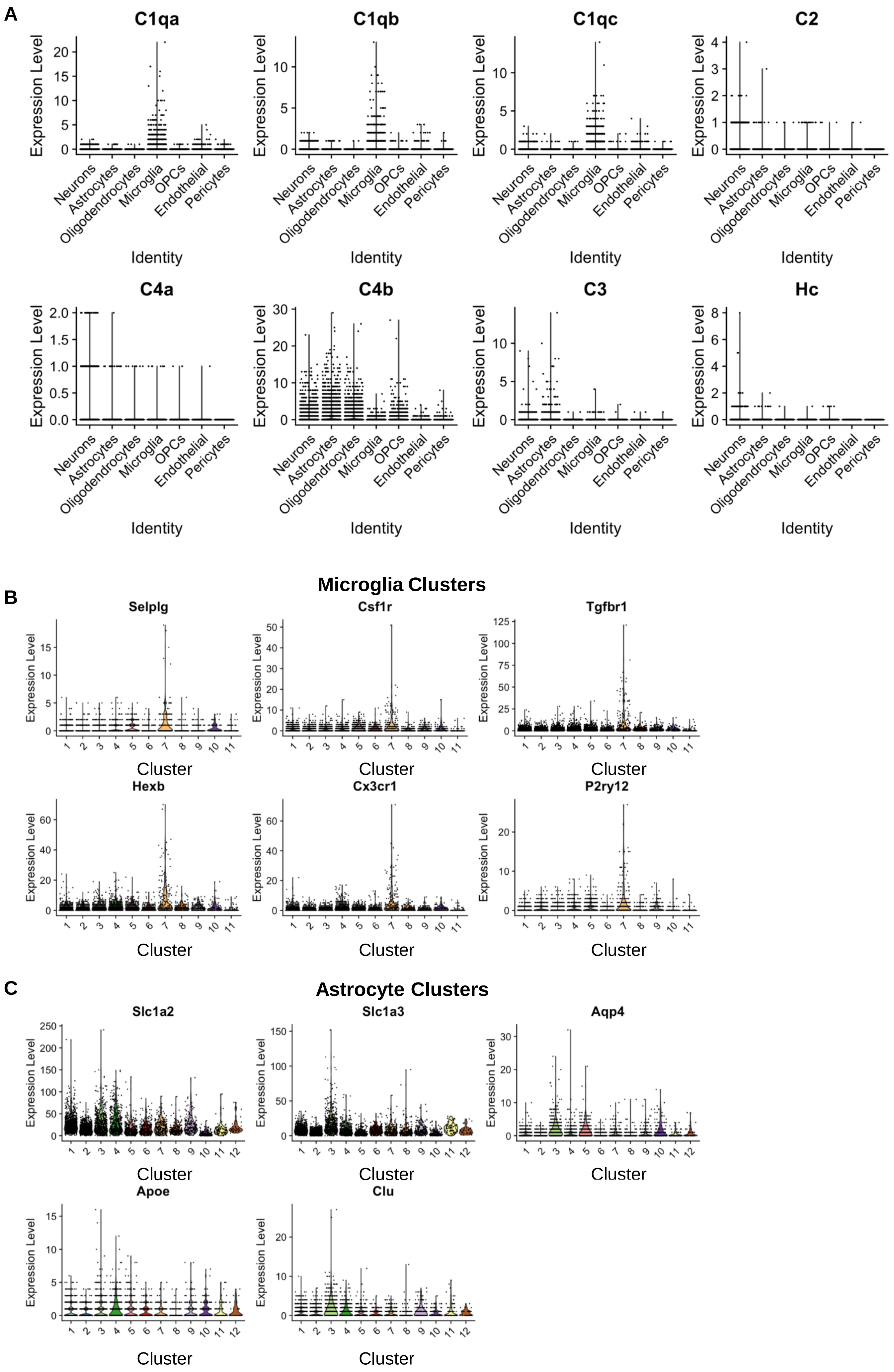
A
C1qa
C1qb
C1qc
C2
C4a
C4b
C3
Hc
Expression Level
Identity
Neurons
Astrocytes
Oligodendrocytes
Microglia
OPCs
Endothelial
Pericytes
Microglia Clusters
B
Selplg
Csf1r
Tgfbr1
Hexb
Cx3cr1
P2ry12
Cluster
Astrocyte Clusters
C
Slc1a2
Slc1a3
Aqp4
Apoe
Clu

**Figure S2: Cell-specific complement expression and validation of microglia and astrocyte clusters with expression of known genes.** (**A**) Expression (count) of complement genes in different cell populations (**B**) Expression levels of microglial genes Selplg, CSF1r, Tgfbr1, Hexb, Cx3cr1, P2ry12 in each microglia cluster. (**C**) Expression levels of astrocytic genes Slc1a2, Slc1a3, Aqp4, Apoe, and Clu in each astrocyte cluster.

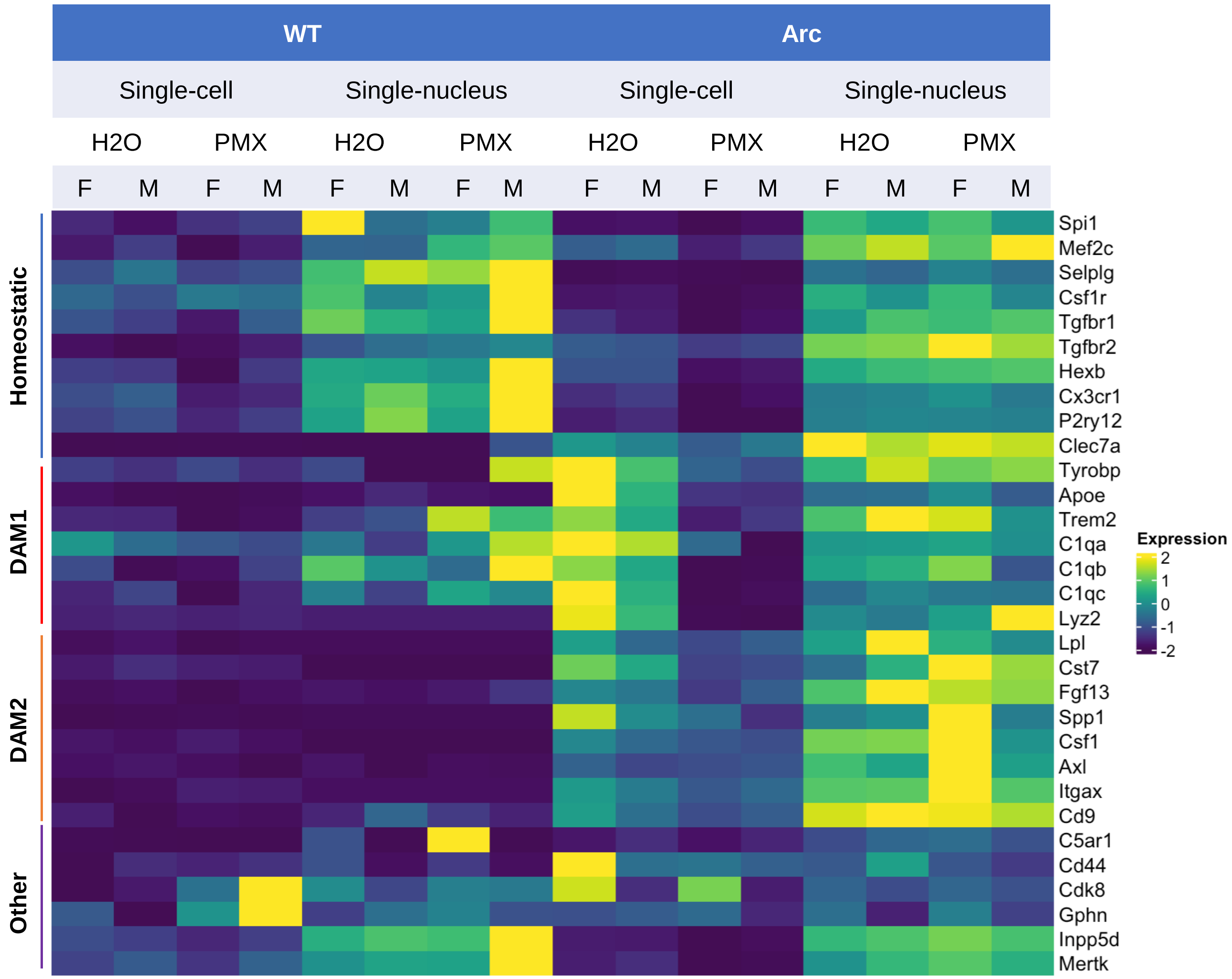


**Figure S3: Comparison of expression of microglial genes in single-cell and single nucleus and by sex.** Relative expression of homeostatic, DAM1, and DAM2 genes was visualized in single-cell and single-nucleus-derived samples. Expression was assessed by genotype, treatment of PMX205, and sex.

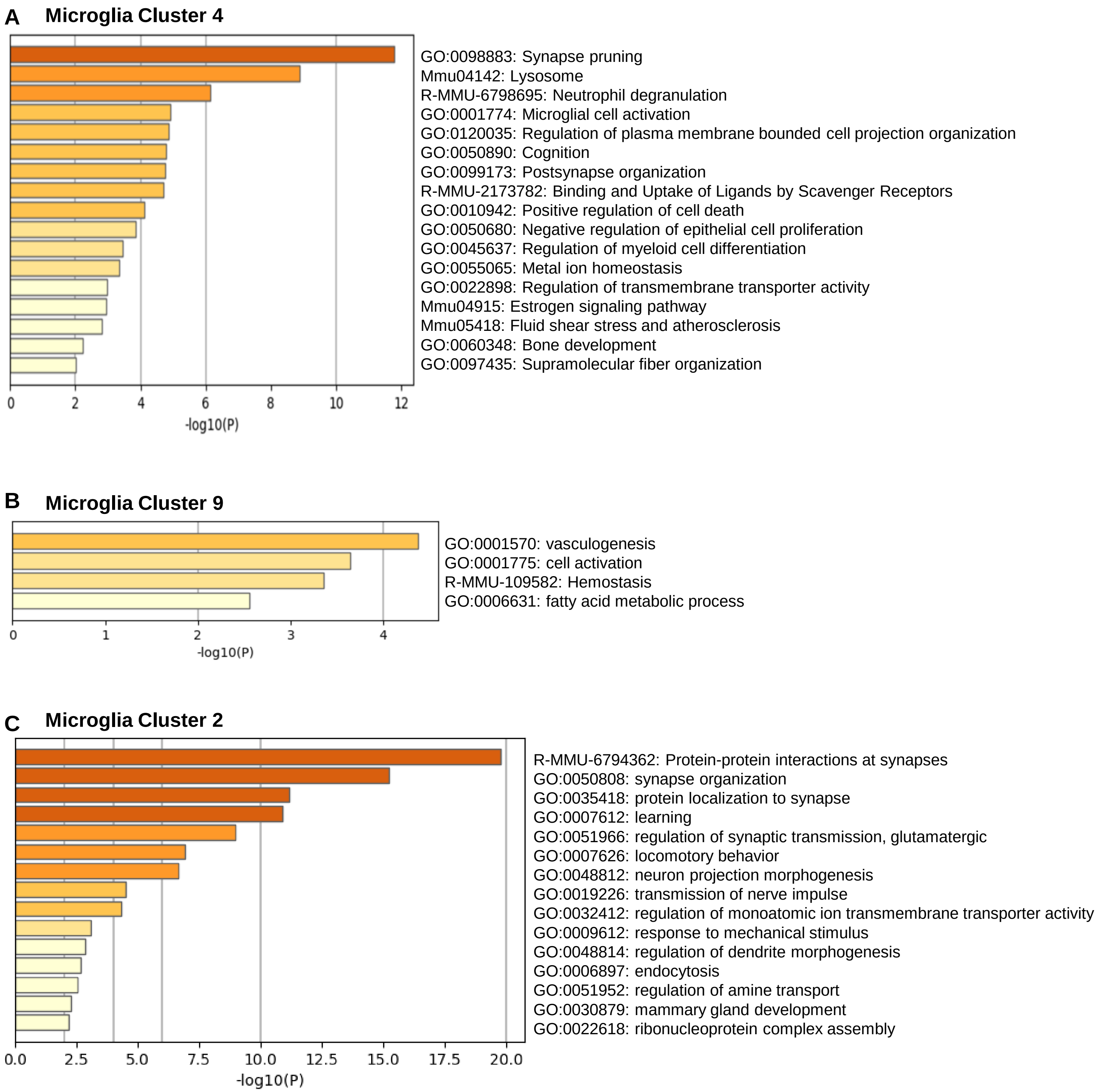


**Figure S4: Gene Ontology analysis of microglia clusters:** Metascape was used to elucidate Gene Ontology (GO) terms for microglial clusters 4 (**A**), 9 (**B**), and 2 (**C**).

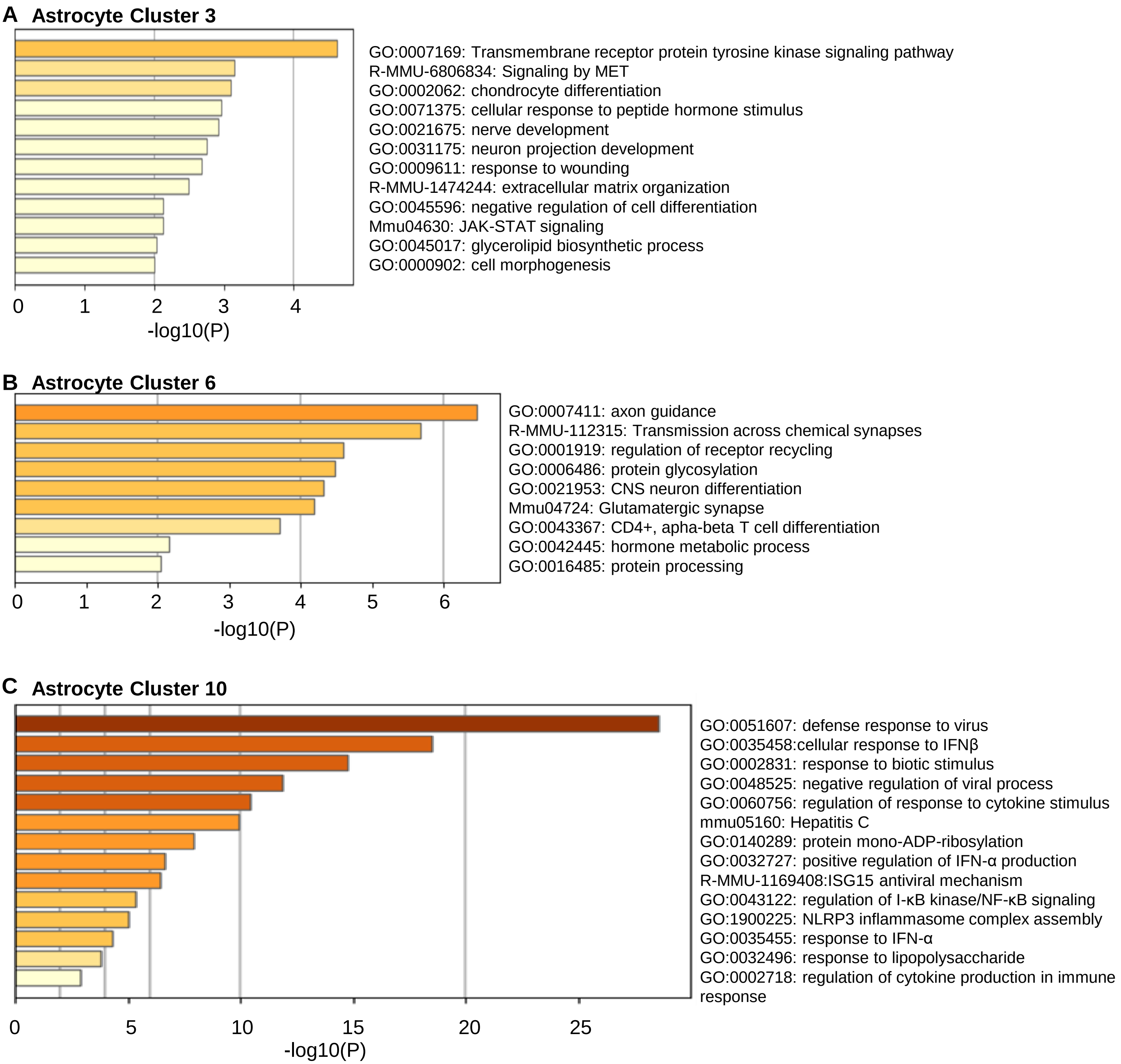


**Figure S5: Gene Ontology analysis of astrocyte clusters:** Metascape was used to elucidate Gene Ontology (GO) terms for microglial clusters 3 (**A**), 6 (**B**), and 10 (**C**).

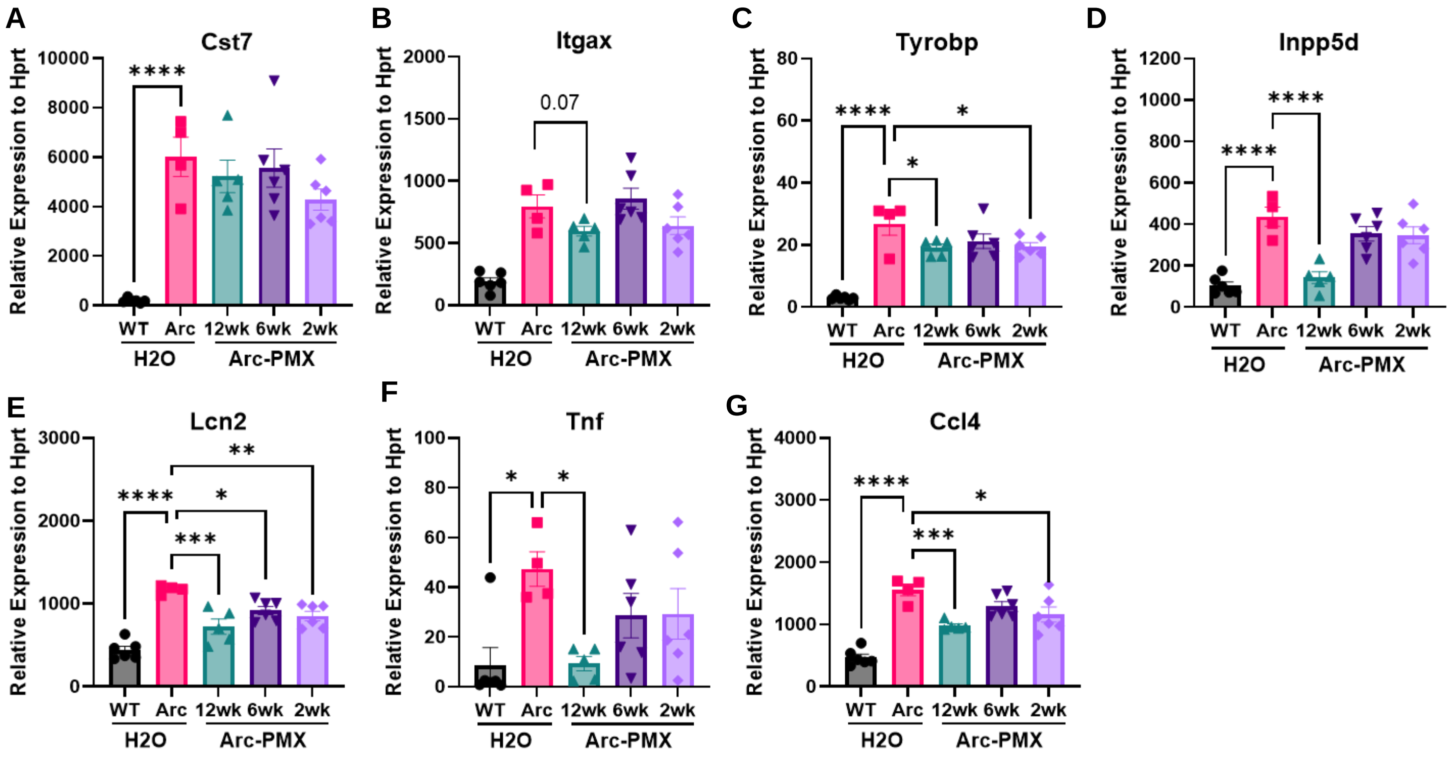


**Figure S6: Reduction in inflammatory gene expression with PMX205 treatment in a younger cohort.** RNA was extracted from hippocampal homogenates derived from 7-month-old mice treated with PMX205 for different durations and analyzed by qPCR for select reactive microglia genes **(A-D)**, reactive astrocyte gene (**E**), or inflammatory cytokines (**F-G**). Data shown as mean ± SEM. One-way ANOVA with Dunnett's *post hoc*. N = 4-6 mice/treatment

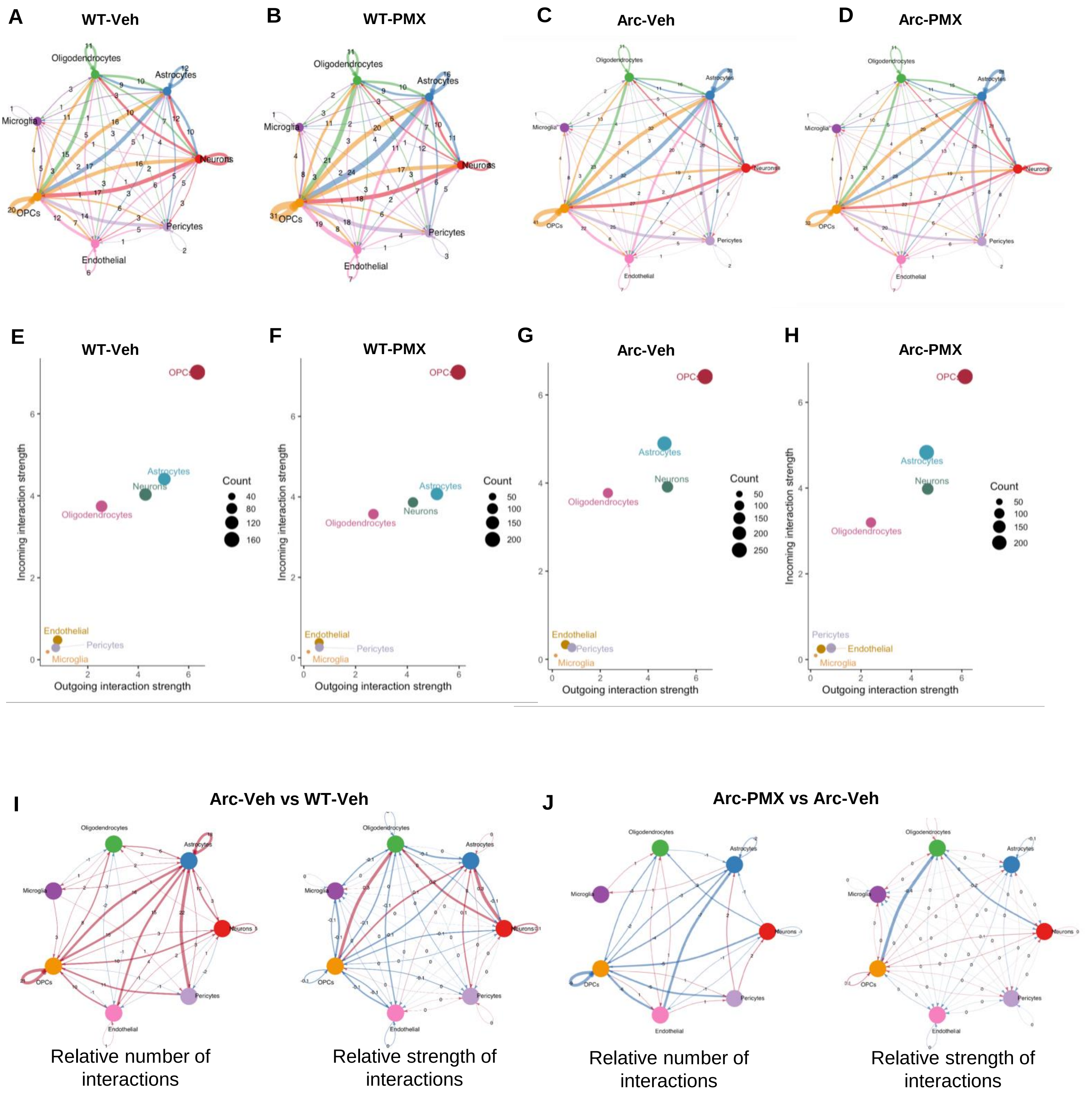


**Figure S7: Aberrant intercellular signaling in Arctic hippocampi is suppressed by C5aR1 inhibition.** (**A-D**) Number of cell-cell interactions in WT-Veh (**A**), WT-PMX (**B**), Arc-Veh (**C**), Arc-PMX (**D**). Cell-specific source and receiver of intercellular signaling shown for WT-Veh (**E**), WT-PMX (**F**), Arc-Veh (**G**), Arc-PMX (**H**). (**I**) Relative number and strength of interactions comparing Arc-veh with WT-veh. (**J**) Relative number and strength of interactions comparing Arc-PMX with Arc-veh.

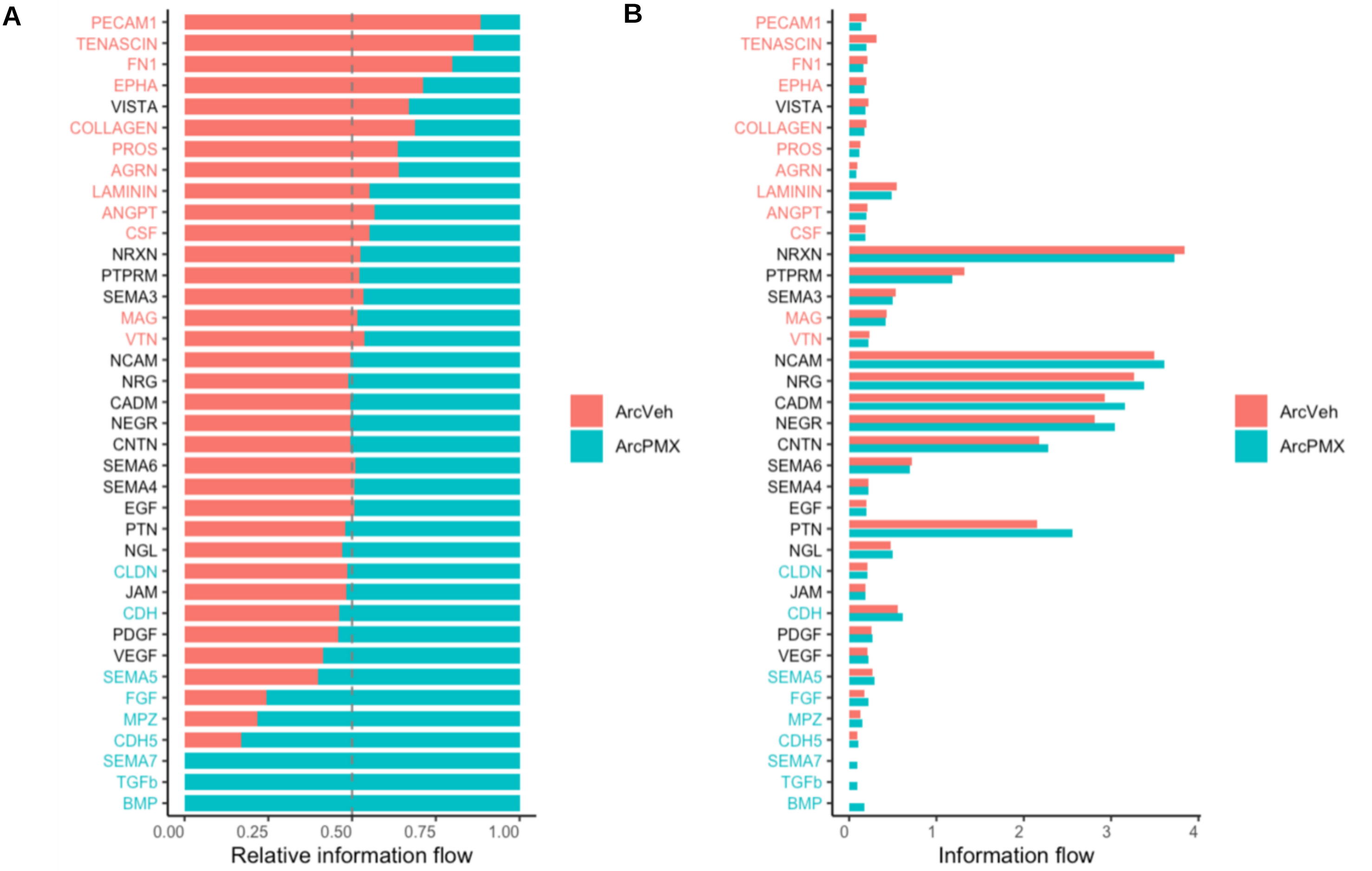


**Figure S8: Relative and absolute information flow of pathways enhanced or suppressed by PMX205 treatment in Arctic mice.** Total list of cell signaling pathways identified by CellChat in Arc-veh vs Arc-PMX. (**A**) Relative information flow of pathways enhanced by PMX205 treatment (text shown in teal), suppressed by PMX205 (text shown in salmon), or unchanged by treatment (text shown in black). (**B**) Absolute information flow of the pathways identified by CellChat in Arc-Veh and Arc-PMX cells.

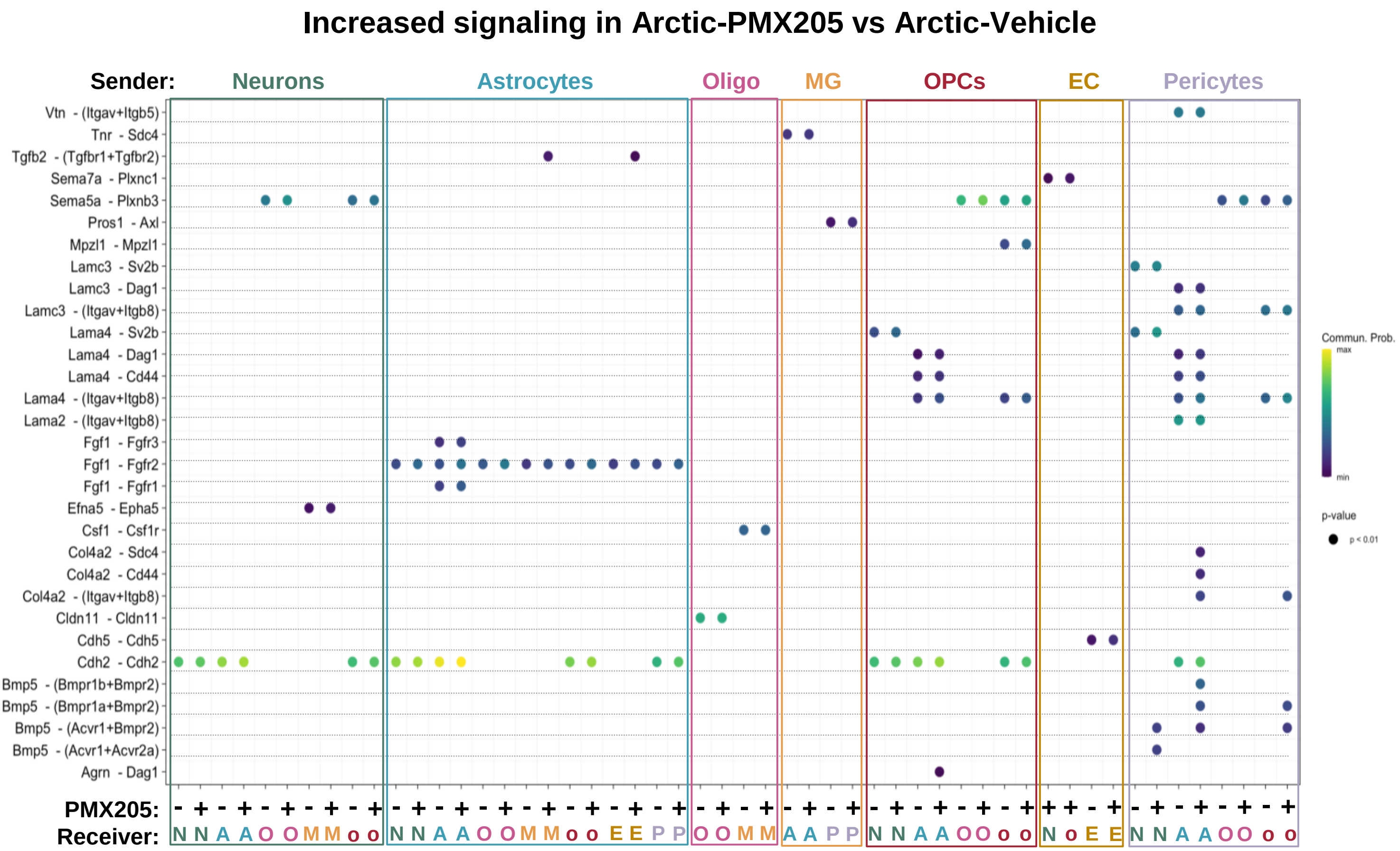


**Figure S9: Receptor-Ligand interactions mediated by different cell types in Arc-Veh and Arc-PMX.** (**A**) Signaling probability of receptor-ligand communication, and cellular sender and receiver in pathways that were increased in Arc-PMX compared to Arc-Veh. (Abbreviations: N, Neuron; A, Astrocyte; O, Oligodendrocyte; M, Microglia; o, OPCs; E, Endothelial Cells; P, Pericytes)

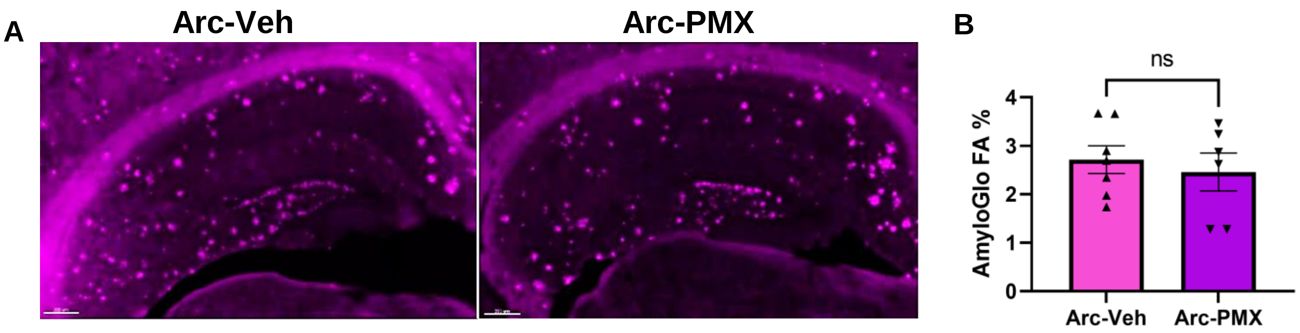


**Figure S10: C5aR1 inhibition does not alter hippocampal plaque deposition in Arctic mice.** (**A**) Representative images of AmyloGlo staining in dorsal hippocampal sections derived from mice 10 months of age. (**B**) Quantification of percent field area of AmyloGlo staining in the hippocampus. Data shown as mean ± SEM. t-test. N = 6-7 mice/genotype/treatment.
